# TSHR-targeting nucleic acid aptamer treats Graves’ ophthalmopathy via novel allosteric inhibition

**DOI:** 10.1101/2025.03.07.641992

**Authors:** Yanchen Zhang, Ende Wu, Weibin Liu, Ling Zeng, Neng Ling, Hongmei Wang, Zhixing Li, Shuang Yao, Tonghe Pan, Xuanwen Li, Yate Huang, Xiaojing Li, Yunhai Tu, Wentao Yan, Jianzhang Wu, Mao Ye, Wencan Wu

**Author notes:** Corresponding author. Email: Jianzhang Wu,; Mao Ye,; Wencan Wu,. These authors contributed equally to this work (Yanchen Zhang, Ende Wu, Weibin Liu).

## Abstract

Abnormal thyrotropin receptor (TSHR) signaling drives a spectrum of diseases affecting hundreds of millions globally, including Graves’ disease (GD), Graves’ ophthalmopathy (GO), hyperthyroidism, and thyroid cancers. Therefore, TSHR is considered an attractive therapeutic target with immense drug discovery potential. Currently, there are no clinically available TSHR inhibitors. To address this unmet medical need, we developed YC3, a functional nucleic acid aptamer targeting TSHR, via an innovative approach combining protein-targeting cell-SELEX and functional selection. YC3 exhibited nanomolar affinity with potent TSHR inhibition. Its therapeutic potential was comprehensively evaluated both *in vitro* and *in vivo* for GO. In GO patient-derived orbital fibroblasts (OFs), YC3 reversed thyroid-stimulating antibodies (TSAbs)-induced cell activation, suppressing inflammatory cytokines and extracellular matrix (ECM) secretion. In a GO mouse model, YC3 treatment markedly attenuated orbital inflammation, ECM deposition, and fibrosis, ameliorating the pathological remodeling of orbital tissue. Mechanistically, YC3 bound to a previously unidentified allosteric site within the leucine-rich repeat domain of TSHR, thereby inhibiting receptor activation. This study identifies YC3 as a promising TSHR-targeted therapy and unveils a new druggable site for inhibitor design. It also provides the first preclinical evidence supporting pharmacological TSHR inhibition for GO treatment, advancing therapeutic development for TSHR-related diseases.

## Introduction

The thyrotropin receptor (TSHR), a pivotal member of the G protein - coupled receptor (GPCR) family, is primarily expressed in thyroid tissue and serves as the central regulator of thyroid physiology. Activated by its endogenous ligand thyrotropin (TSH), TSHR initiates signaling cascades that drive thyroid hormone synthesis and secretion, which are essential for metabolic homeostasis^1, 2^. Beyond its canonical thyroidal role, TSHR can be found in extrathyroidal tissues such as the brain, thymus, and orbit, where it mediates immune regulation and neurobehavioral processes ^3, 4^. Aberrant TSHR signaling has been involved in the pathogenesis and progression of various disorders, including Graves’ disease, Graves’ ophthalmopathy (GO), hypothyroidism, thyroid cancer, and certain neurological disorders ^4, 5^, affecting hundreds of millions worldwide. Among these disorders, GO, also known as thyroid-associated ophthalmopathy (TAO), is an autoimmune disorder affecting the orbit. It is characterized by proptosis, eyelid retraction, and restrictive strabismus, which can lead to disfigurement, vision loss, and a reduced quality of life ^6, 7^. Currently, intravenous methylprednisolone is the primary treatment for GO, but it has limited efficacy and can cause serious side effects ^8^. Given these limitations, the clinical needs are unmet, highlighting the urgent requirement for more effective treatment strategies for GO. Moreover, the expression of TSHR is significantly upregulated in virtually all orbital tissues affected by GO ^9–12^, and thyroid-stimulating antibodies (TSAbs) targeting TSHR are prevalent in GO patients. These TSAbs correlate strongly with GO’s clinical activity and severity^13–15^. The availability of TSHR immunization-induced GO animal models^16^, combined with GO’s organ-specific disorder that allows for localized drug administration, provides an ideal preclinical platform for evaluating TSHR-targeted therapeutics *in vivo*.

The significant clinical need for TSHR inhibitors makes TSHR an attractive target for drug discovery and development ^4^. Over the past decade, various TSHR inhibitors such as small molecules, monoclonal antibodies, and peptides have emerged^4, 17, 18^. However, most exhibit suboptimal characteristics like micromolar potency, insufficient selectivity, and poor pharmacological properties. Only the binding mechanism of the orthosteric inhibitor K1-70 monoclonal antibody has been elucidated ^2^, leaving the molecular interactions of other inhibitors undefined. This knowledge gap, along with the lack of clinically approved TSHR inhibitors, underscores the urgent need to develop mechanically well-characterized and effective TSHR inhibitors.

Aptamers are an innovative class of therapeutic molecules consisting of single-stranded nucleic acids, typically 20 to 100 nucleotides in length. They form a unique three-dimensional structure that enables them to bind with high affinity and specificity to their target ^19^. These molecules offer distinct advantages over conventional therapeutics, including ease of synthesis and modification, low toxicity, cost-effectiveness, and low immunogenicity^20, 21^. The clinical potential of aptamers is highlighted by FDA-approved therapies like Pegaptanib and Zimura ^22^.

In this study, we employed an innovative tandem approach combining protein-targeted cell-SELEX with specific functional selection to develop YC3, a TSHR-targeting inhibitory aptamer. Our findings indicate that YC3 exerts its inhibitory effects by binding to a previously unidentified allosteric site on the TSHR. This novel mechanism demonstrates YC3’s therapeutic potential for GO, supported by robust biological activity in both *in vitro* and *in vivo* systems.

## Results

### Subhead 1: Screening and identification of TSHR-targeting aptamers with inhibitory function

To screen TSHR-targeting aptamers with inhibitory function, we employed a tandem process combining protein-targeted cell-SELEX and a specific functional selection process (Fig. 1A). For protein-targeted cell-SELEX, HEK293T cells stably expressing TSHR (TSHR-293T) were used as positive cells, while HEK293T cells transfected with an empty vector (MOCK) served as negative controls (Fig. S1A). The expression levels of TSHR mRNA and protein were high in the TSHR-293T cells, but not high in the MOCK cells detected by qPCR, flow cytometry and western blotting (Fig. S1B-D). Subsequently, cell-SELEX was conducted using TSHR-293T cells for positive selection and MOCK cells for negative selection. The enrichment of the ssDNA sequences in different rounds was monitored by flow cytometry. With each subsequent screening round, the fluorescence intensity of the TSHR-293T cells progressively increased, whereas that of the MOCK cells remained stable. The maximum fluorescence intensity of the screening round was observed in round 9 (Fig. 1B). Therefore, the ssDNAs enriched in round 9 were selected for high-throughput sequencing using Illumina MiSeq.

**Fig. 1.**
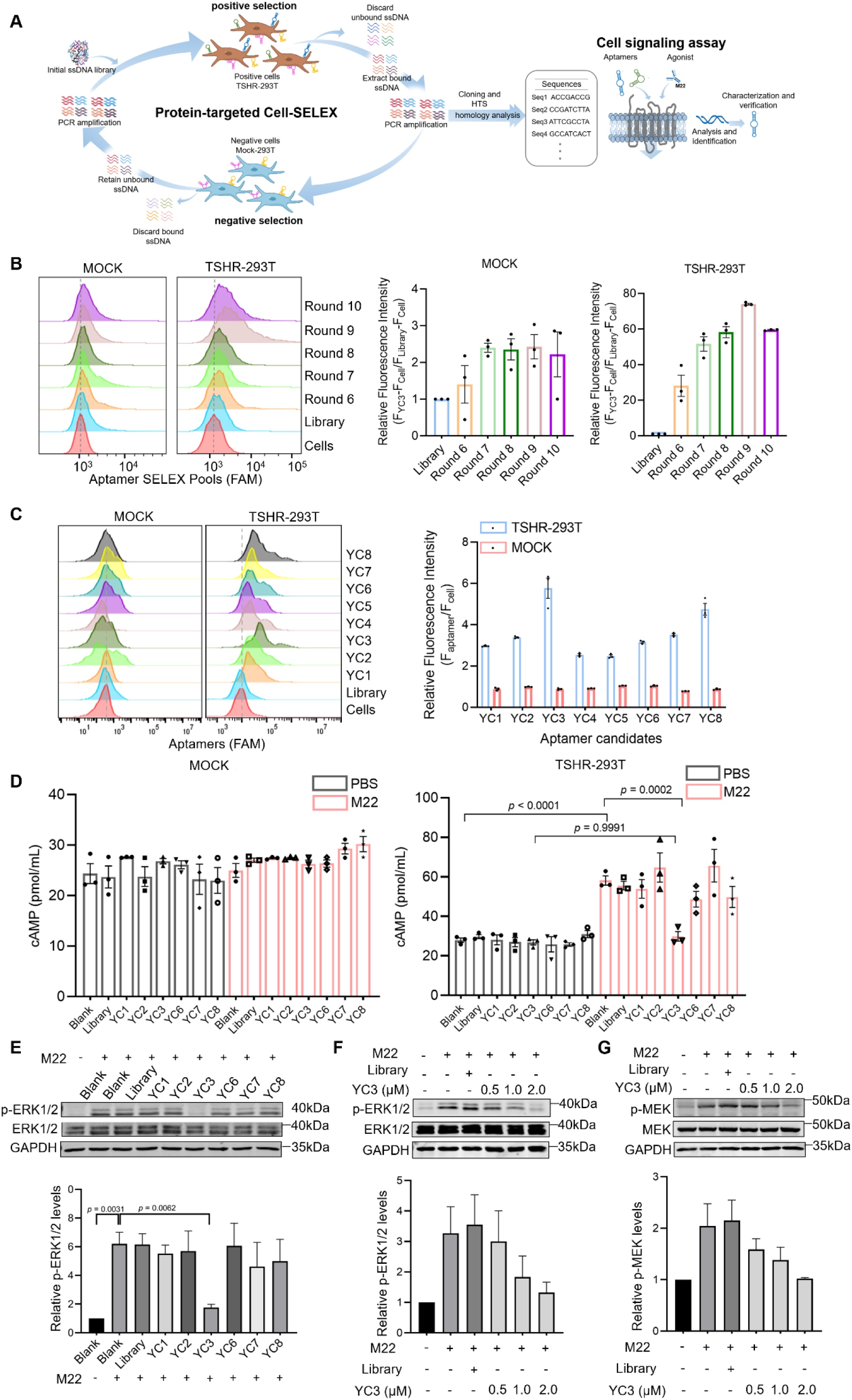
Screening of TSHR-targeting DNA aptamers with inhibitory function. (**A**) Schematic illustration of aptamer inhibitors selection strategy for TSHR. (**B**) Binding assays of the selected pool with MOCK and TSHR-293T cells by flow cytometry. The FAM-labeled initial Library was used as the control. Quantitative analysis of the relative fluorescence intensity of the aptamer candidates in TSHR-293T and MOCK cells, n=3 independent samples in each group. Data are represented as mean±SEM. (**C**) The binding of candidate aptamers (250 nM) to MOCK and TSHR -293T cells was analyzed by flow cytometry. Quantitative analysis of the relative fluorescence intensity of the aptamer candidates in TSHR-293T and MOCK cells, n=3 independent samples in each group. Data are represented as mean±SEM. (**D**) The intracellular cAMP levels in MOCK and TSHR-293T cells, treated with or without aptamer candidates and M22, were measured by ELISA, n=3 independent samples in each group. Data are represented as mean±SEM. One-way ANOVA, followed by Tukey’s multiple post hoc test was used to calculate *P* values. (**E**) Representative immunoblots/densitometric quantitative analysis of phospho-ERK1/2 protein level and ERK1/2 protein level in TSHR-293T cells stimulated by M22 and treated with aptamer candidates. GAPDH was used as a loading control, n=3 independent samples. Data are represented as mean±SEM. Statistics: one-way ANOVA followed by Tukey’s multiple post hoc test. (**F**) Representative immunoblots/densitometric quantitative analysis of phospho-ERK1/2 protein level and ERK1/2 protein level in TSHR-293T cells stimulated by M22 and treated with series concentrations of YC3. GAPDH was used as a loading control, n=3 independent samples in each group. Data are represented as mean±SEM. (**G**) Representative immunoblots/densitometric quantitative analysis of phospho-MEK protein level and MEK protein level in TSHR-293T cells stimulated by M22 and treated with series concentrations of YC3. GAPDH was used as a loading control, n=3 independent samples in each group. Data are represented as mean±SEM.

Based on abundance and homogeneity, the sequenced ssDNAs were classified into six families (Fig. S2). Eight representative sequences (YC1 to YC8) were synthesized as aptamer candidates for further evaluation. The binding ability of these aptamers was assessed by flow cytometry. Interestingly, all eight sequences showed specific binding to the TSHR-293T cells but not to the MOCK cells. Among them, YC1, YC2, YC3, YC6, YC7, and YC8 exhibited relatively intense fluorescence signals (Fig. 1C). Generally, ligand binding induces conformational changes in TSHR, leading to the activation of multiple intracellular signaling pathways. Previous studies have shown that activating TSHR with M22, a human TSAb, increases intracellular cyclic adenosine monophosphate (cAMP) production and the phosphorylation of ERK1/2^4, 23^. To further screen inhibitory aptamers, a specific functional selection process based on cAMP level and p-ERK1/2 expression was employed. Notably, under non-stimulated conditions, none of the aptamer candidates influenced cAMP production in either MOCK cells or TSHR-293T cells. In contrast, YC3 markedly inhibited cAMP production in M22-stimulated TSHR-293T cells (Fig. 1D). Furthermore, YC3 effectively downregulated M22-induced phosphorylation of ERK1/2 in TSHR-293T cells (Fig. 1E). Immunoblotting showed the inhibitory effect of YC3 on M22-induced phosphorylation of both ERK1/2 and MEK in a dose-dependent manner (Fig. 1F-G). These findings collectively demonstrate that YC3 has significant potential to inhibit TSHR downstream signaling activation. Based on its specific binding and inhibitory effects, YC3 was selected for further investigation.

### Subhead 2: Optimization, characterization, and modification of aptamer YC3

In general, aptamers specifically bind to targets through unique three-dimensional structures determined by sequences. To further optimize the sequence of YC3, we designed two truncated schemes based on the predicted secondary structure using NUPACK (**{**HYPERLINK “https://www.nupack.org/”**}**). The full-length YC3 was truncated by removing the bases that formed the terminal stem-loop structures (Fig. S3A). The resulting sequences are presented in Table S1. Flow cytometry showed that truncated YC3 sequences diminished binding ability to TSHR-293T cells, highlighting the importance of the full YC3 structure for high affinity (Fig. S3B). Additionally, flow cytometry revealed that the equilibrium dissociation constant (K_d_) of YC3 against TSHR-293T cells was 243.2 ±54.5 nM (Fig. 2B), indicating a strong binding affinity between YC3 and TSHR-293T cells. It has been reported that the structures of aptamers can change slightly at different temperatures, which may affect their affinity^24^. Since YC3 was generated using cell-SELEX at 4°C, we investigated its binding ability at physiological temperature. Incubation of YC3 with TSHR-293T cells at 37°C produced a stronger fluorescence signal compared to incubation at 4°C (Fig. 2C), indicating that physiological temperature enhances the binding affinity of YC3.

**Fig. 2.**
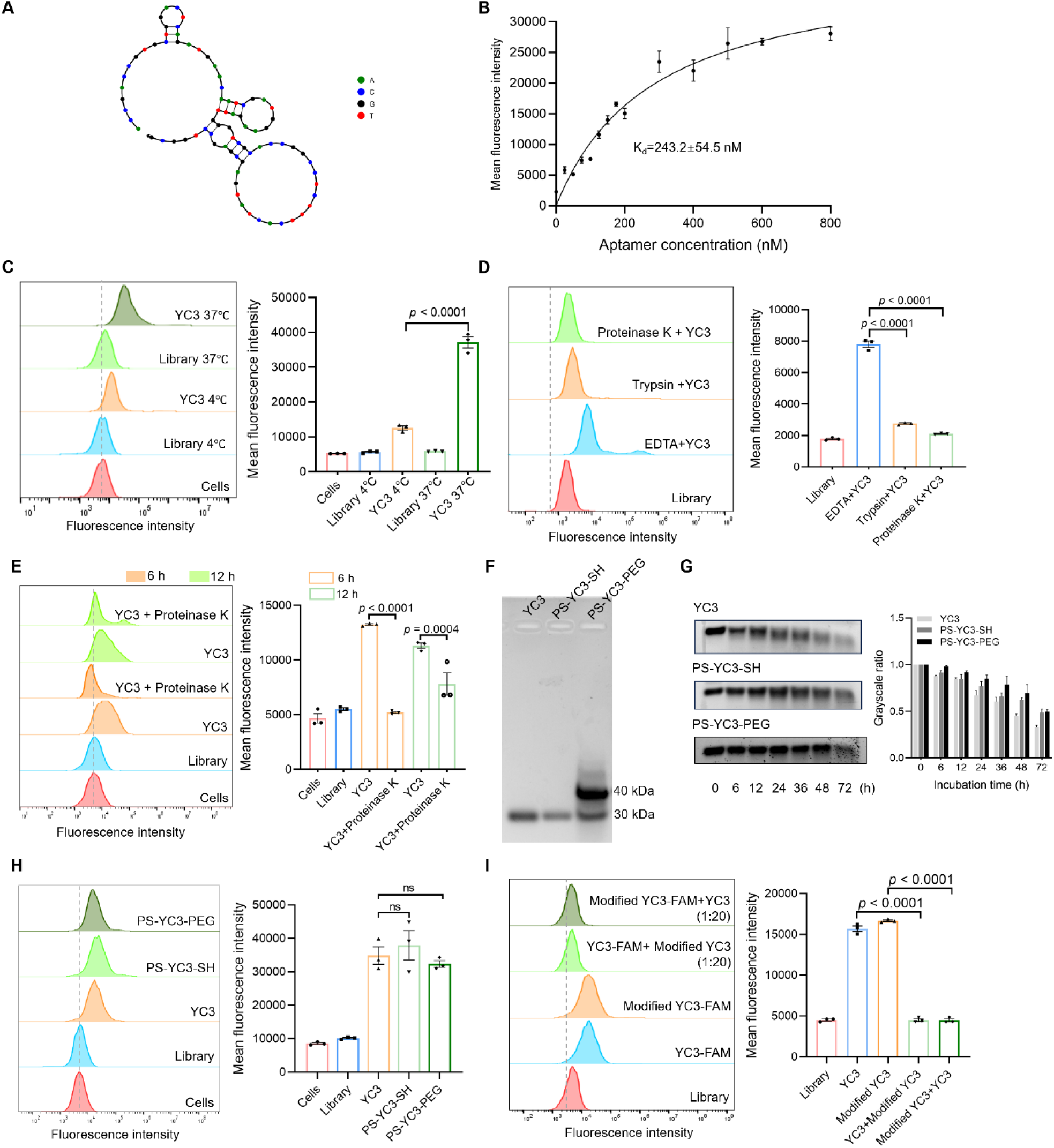
Characterization and modification of aptamer YC3. (**A**) The secondary structure of YC3 was predicted by NUPACK software. (**B**) The dissociation constant of YC3 for TSHR-293T cells was determined by flow cytometry, using the following equation: Y=B_max_ X/(K_d_+X) (X representing the concentration of aptamer, Y representing the mean fluorescence intensity, and B_max_ representing the peak fluorescence intensity), n=3 independent samples. (**C**) Flow cytometry was employed to determine the binding ability of YC3 for TSHR-293T cells at 4°C or 37°C. The FAM-labeled initial Library was used as the control. Quantitative analysis of mean fluorescence intensity of YC3, n=3 independent samples in each group. Data are represented as mean±SEM. Two-tailed Student’s t-test was used to calculate *P* values. (**D**) Flow cytometry was employed to determine the binding ability of YC3 for TSHR-293T cells treated with 0.25% trypsin or 10 µg/mL proteinase K. The TSHR-293T cells treated with the EDTA were used as the control. Quantitative analysis of mean fluorescence intensity of YC3, n=3 independent samples in each group. Data are represented as mean±SEM. One-way ANOVA, followed by Tukey’s multiple post hoc test was used to calculate *P* values. (**E**) Flow cytometry was employed to determine the binding ability of YC3 for TSHR-293T cells for 6 hours or 12 hours with or without treatment of 10 µg/mL proteinase K. Quantitative analysis of mean fluorescence intensity of YC3, n=3 independent samples in each group. Data are represented as mean±SEM. One-way ANOVA, followed by Tukey’s multiple post hoc test was used to calculate *P* values. (**F**) Gel electrophoresis analysis of unmodified YC3 and its two modified forms. PS-YC3-SH involves replacing the standard oligonucleotides at both termini with phosphorothioate oligonucleotides and adding a sulfhydryl group to the 3’ end, without undergoing PEGylation. PS-YC3-PEG follows the same modification but includes an additional PEGylation step. (**G**) Representative agarose gel electrophoresis /densitometric quantitative analysis of serum stability of YC3 and its modified forms in mouse serum, n=3 independent samples in each group. Data are represented as mean±SEM. (**H**) Flow cytometry was employed to compare the binding ability between YC3 and its modified forms, n=3 independent samples in each group. Data are represented as mean±SEM. One-way ANOVA, followed by Tukey’s multiple post hoc test was used to calculate *P* values. (**I**) Flow cytometry was employed to analyze the competitive binding between YC3 and PS -YC3-PEG (Modified YC3) with indicated concentrations, n=3 independent samples in each group. Data are represented as mean±SEM. One-way ANOVA, followed by Tukey’s multiple post hoc test was used to calculate *P* values.

To assess the effect of cell membrane protein removal on YC3 binding, TSHR-293T cells were treated with trypsin or proteinase K before incubation with YC3. Treatment with either enzyme significantly reduced YC3 binding to TSHR-293T cells (Fig. 2D), indicating that YC3 targets are located on the cell membrane, aligning with the known membrane location of TSHR. To assess the internalization of YC3, TSHR-293T cells were incubated with FAM-labeled YC3 for 6 and 12 hours at 37°C. Post-incubation, proteinase K was applied to remove any YC3 remaining on the cell surface. Flow cytometry analysis revealed a significant reduction in fluorescence intensity after proteinase K treatment, indicating that YC3 did not internalize into TSHR-293T cells within the 12-hour incubation period (Fig. 2E).

Aptamers are widely known for their poor stability and high elimination rate *in vivo*. Therefore, enhancing the biological stability of aptamers is critical for both *in vivo* and *in vitro* applications. To address this, we modified YC3, resulting in PS-YC3-PEG, by substituting five standard oligonucleotides at both termini with phosphorothioate oligonucleotides, adding a sulfhydryl group to the 3’ end, and subsequently performing PEGylation. After modification, the molecular weight of YC3 increased, and its serum stability was enhanced without compromising its binding ability to TSHR-293T cells (Fig. 2F-H). We further investigated the binding sites of YC3 and PS-YC3-PEG through competitive binding analysis. As illustrated in Figure 2I, excess unlabeled PS-YC3-PEG inhibited the binding of FAM-labeled YC3 to TSHR-293T cells, and excess unlabeled YC3 similarly inhibited FAM-labeled PS-YC3-PEG. This suggests that YC3 and PS-YC3-PEG share the same binding sites on TSHR-293T cells.

### Subhead 3: Verification of TSHR as the binding target of YC3

To confirm whether TSHR is the bona fide binding target of YC3, TSHR-specific siRNAs were transfected into Nthy-ori 3-1 cells to knock down TSHR expression. Flow cytometry analysis revealed that TSHR silencing reduced TSHR expression and correspondingly decreased YC3 binding to Nthy-ori 3-1 cells (Fig. 3A). To further investigate the relationship between YC3 binding and TSHR expression across different cell types, we analyzed membrane TSHR expression levels using TSHR-specific antibodies in thyroid, thyroid carcinoma, and ocular cell lines. TSHR was expressed in the thyroid cell line (Nthy-ori 3-1) and thyroid cancer cell lines (BCPAP and FTC-133), but was absent in ocular cell lines (ARPE19, HREC, HLEC, HMC3, and HECT). The binding ability of YC3 to these eight cell lines correlated with membrane TSHR expression levels (Fig. 3B). The Pearson correlation coefficient between TSHR expression levels and YC3 binding levels across cell lines was 0.9251 (Fig. 3C), indicating a strong correlation between YC3 binding ability and TSHR expression levels. To further confirm the specificity of YC3, membrane proteins of TSHR-293T cells were extracted and subjected to an aptamer pull-down assay. As shown in Fig. 3D, TSHR was detected in the protein pulled down by the YC3 group but not in the control Bead group or Library group, demonstrating that YC3 binds specifically to TSHR. Additionally, the binding affinity between the TSHR recombinant protein and the aptamer YC3 was determined by surface plasmon resonance (SPR), yielding a K_d_ of 11.74 nM (Fig. 3E). Taken together, these findings conclusively demonstrate that YC3 specifically binds to the TSHR protein with high affinity, further supporting its potential for TSHR-targeted applications.

**Fig. 3.**
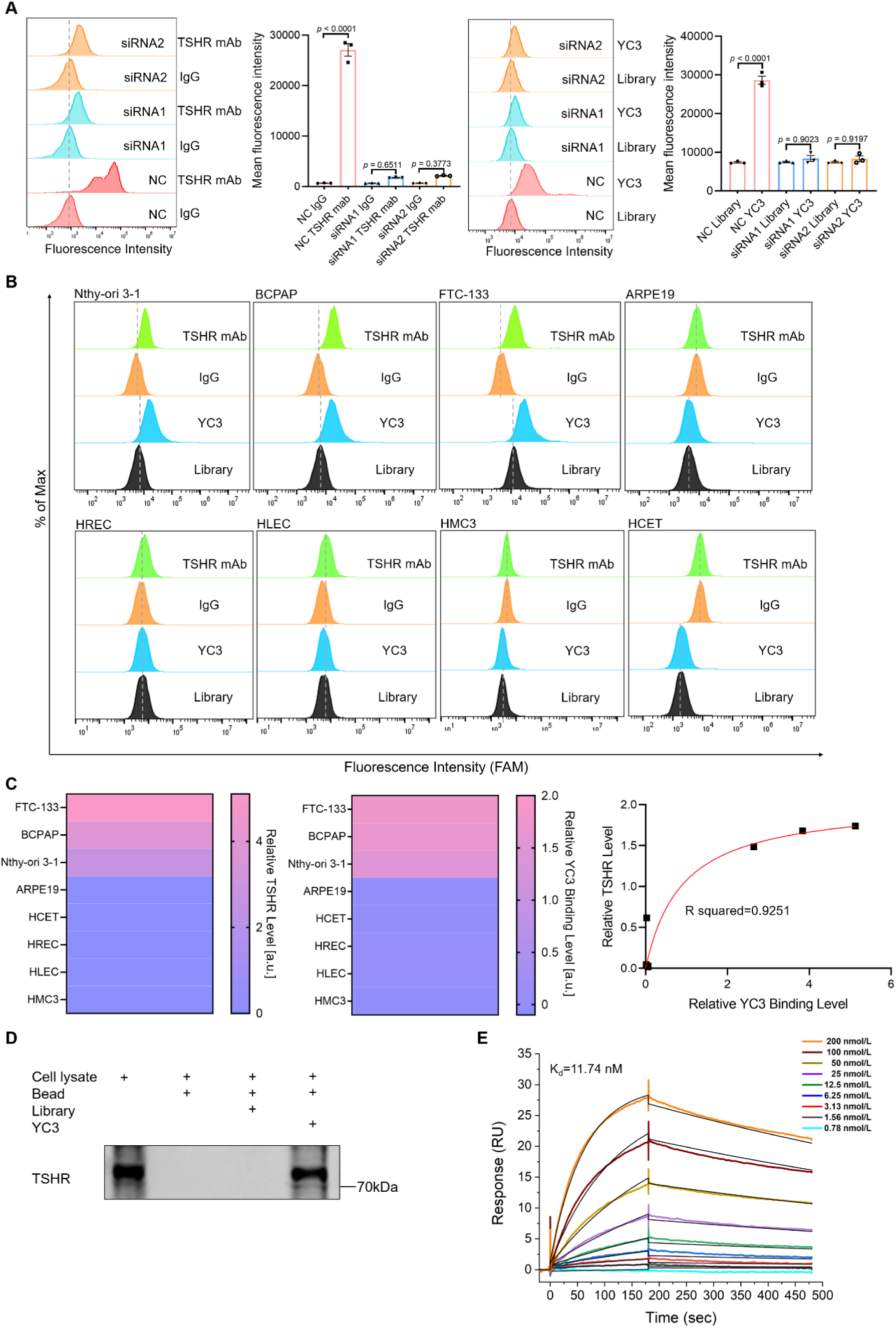
Verification of TSHR as the binding target of aptamer YC3. (**A**) Flow cytometry analysis of TSHR expression levels using TSHR mAb, as well as the binding ability of YC3 to Nthy-ori 3-1 cells following transfection with TSHR siRNA1 or TSHR siRNA2. Quantitative analysis of the relative fluorescence intensity, n=3 independent samples in each group. Data are represented as mean±SEM. One-way ANOVA, followed by Tukey’s multiple post hoc test was used to calculate *P* values. (**B**) Flow cytometry was employed to analyze TSHR protein expression levels using TSHR mAb, with IgG serving as a negative control, and evaluate the binding affinity of aptamer YC3, with an initial pool Library serving as a negative control, in various cell lines including thyroid (Nthy-ori 3-1), thyroid carcinoma (BCPAP and FTC-133), and ocular (ARPE19, HREC, HLEC, HMC3, and HCET) cell lines. (**C**) Heatmap of the relative TSHR levels and YC3 binding levels in different cell lines, along with the correlation between the relative TSHR levels and YC3 binding levels across these cell lines. (**D**) Aptamer pull-down assay. Lane 1: cell lysate; Lane 2: the proteins pulled down by naked beads; Lane 3: the proteins pulled down by aptamer Library-conjugated beads; Lane 4: the proteins pulled down by aptamer YC3-conjugated beads. Samples were stained with TSHR antibody. (**E**) A standard kinetics SPR assay demonstrated the binding of YC3 to immobilized TSHR, with a dissociation constant (K_d_) of 11.74 nM.

### Subhead 4: Identification of the YC3-binding site on the TSHR protein

In humans, the mature full-length TSHR protein is composed of an extracellular region, which includes a large leucine-rich repeat domain (LRD) and a hinge region, a transmembrane domain, and an intracellular region^4^. TSH and TSAbs primarily bind to the concave surface of the receptor’s LRD^2^. To investigate the binding sites between the aptamer YC3 and TSHR, we conducted a series of experiments to characterize the detailed interaction region. Initially, we performed a competitive binding analysis to determine if YC3 competes with M22 for TSHR binding. Flow cytometry showed that neither excess M22 nor YC3 inhibited the other’s binding (Figure 4A-B), which indicates that YC3 and M22 bind to different sites on the TSHR. To define the specific interaction region, we performed molecular dynamics simulations to map the YC3-binding residues on TSHR. In a blind docking process, 100 conformations were generated, clustered, and ranked according to their docking energy values. The 50 conformations with the highest number of clusters and the lowest binding energies were selected to predict the protein-aptamer binding interface. Analysis of these conformations revealed that, unlike M22, YC3 is bound to the convex surface of the TSHR LRD (Fig. S4A). The complex structure with the lowest binding score from the precise docking is presented in Figure 4C-E. To characterize the dynamic behavior of the TSHR-YC3 complex, a 100 ns molecular dynamics simulation was performed, and the refined binding model was extracted from the MD trajectory. The binding interface is primarily located in two regions (Fig. 4F), designated A and B, with the interaction dominated by a robust hydrogen bond network (Fig. S4B). Key residues of TSHR that form important interactions with YC3 include T190, Q193, Q173, N170, K146, L265, L266, S268, S243, K244, K218, D219, Y195, and N198 (Fig. 4G-H). Residue binding free energy decomposition analysis revealed that K218, D219, K244, and N170 made the largest contributions to the binding energy of the complex in both the docking stage and the kinetic simulation (Table S2).

**Fig. 4.**
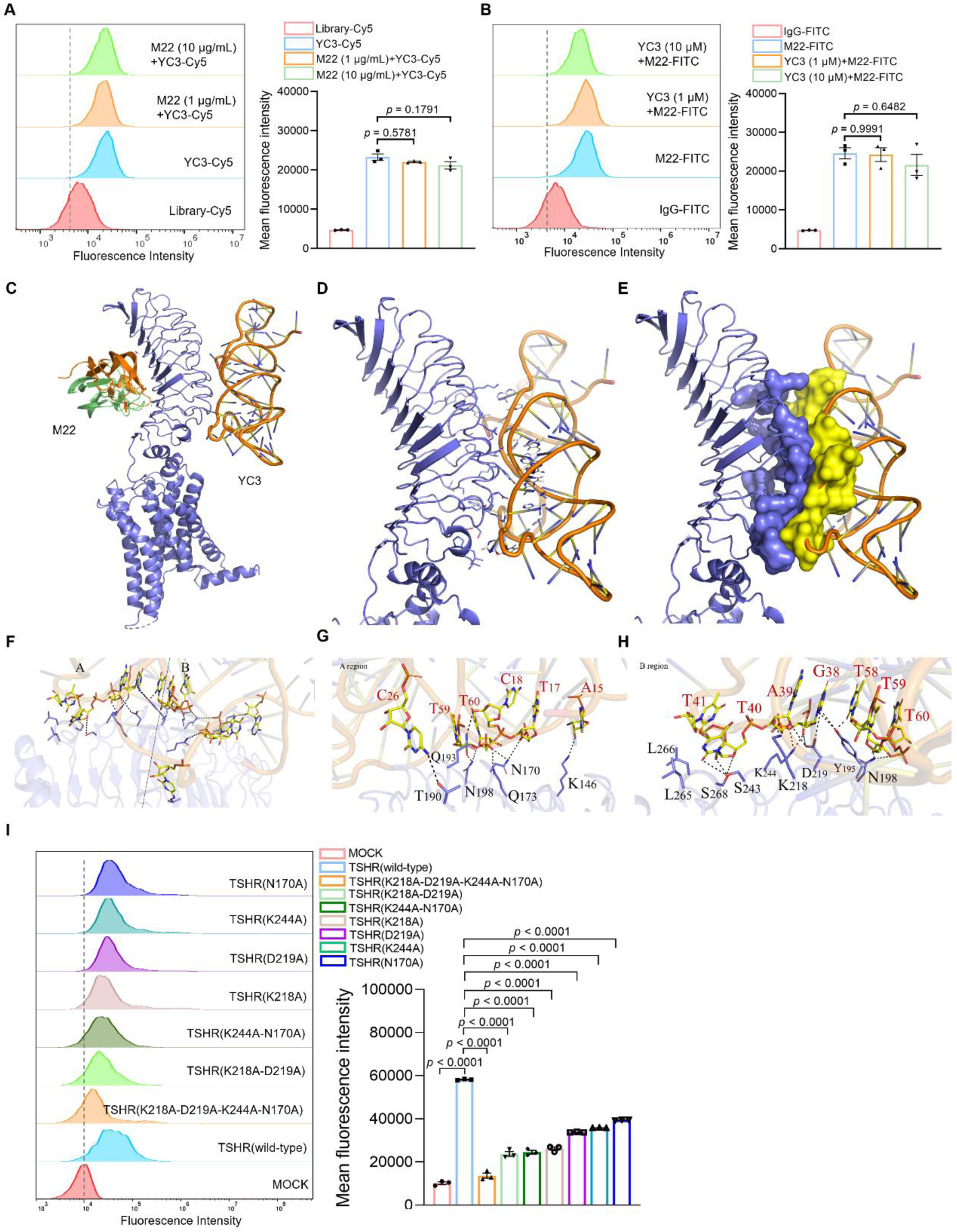
Identification of the binding site of aptamer YC3 on the TSHR protein. (**A** and **B**) Competitive binding assay of YC3 and M22 by flow cytometry. Quantitative analysis of the relative fluorescence intensity, n=3 independent samples in each group. Data are represented as mean±SEM. One-way ANOVA, followed by Tukey’s multiple post hoc test was used to calculate *P* values. (**C**) Molecular dynamic simulation results of binding sites at the interface between aptamer YC3, M22 and TSHR. (**D** and **E**) Close view of TSHR and YC3 interaction. (**F**) Two regions (A region shown in **G** and B region shown in **H**) of the binding sites between aptamer YC3 and TSHR were enlarged and represented as stick models. The key residues are marked in black. (**I**) Flow cytometry results of binding between YC3 and HEK293T cells expressing wild-type and mutant TSHR proteins. Quantitative analysis of the relative fluorescence intensity, n=3 independent samples in each group. Data are represented as mean±SEM. One-way ANOVA, followed by Dunnett’s multiple comparisons test was used to calculate *P* values.

To validate computationally predicted binding residues, we performed site-directed mutagenesis on four key residues, mutating them to Alanine in quadruple, double, and single mutants. Flow cytometry assessed YC3 binding to HEK293T cells expressing these TSHR mutations. TSHR expression levels were comparable across all transfected cell types (Fig. S9). As shown in Fig. 4I and Fig. S9, YC3 binding was most significantly reduced in cells with quadruple mutants, followed by double mutants, and then single mutants. The K218A single mutant caused the most substantial decrease in YC3 binding, highlighting its critical role. These findings support the molecular dynamics simulations, confirming the importance of these residues for YC3 binding. In summary, our results provide molecular insights into the TSHR binding site for YC3, revealing a distinct epitope on the convex surface of the LRD. The identification of key interacting residues suggests a potential mechanism for allosteric modulation of TSHR by YC3.

### Subhead 5: Aptamer YC3 targeting orbit in vivo and in vitro

TSHR is considered the primary autoantigen in GO, and OFs are the key effector cells and targets of autoimmune attacks^25^. In GO development, enhanced TSHR expression is driven by autoimmune and inflammatory processes, paralleling de novo adipogenesis^3^. Compared to healthy donors, TSHR expression was significantly higher in orbital tissues from GO patients (Fig. S5A-B). We then investigated YC3 binding in orbital cells and tissues. Graves’ orbital fibroblasts (GOFs) and normal orbital fibroblasts (NOFs) were isolated from GO patients and healthy donors, respectively, and identified using the fibroblast marker Vimentin (Fig. S5C). YC3 showed strong binding affinity to GOFs, with a K_d_ of 99.23 ± 29.8 nM (Figure 5B). Oil Red O staining revealed that fibroblasts with adipogenic differentiation became rounded and accumulated red-stained lipid droplets (Fig. S5D). Notably, adipogenic differentiation significantly increased TSHR expression in GOFs but not in NOFs (Fig. S5E-F). Given these differences, we examined YC3 binding to GOFs and NOFs in both undifferentiated and adipogenic states. YC3 binding was stronger in GOFs than in NOFs, and even stronger in adipogenically differentiated GOFs (Figure 5C).

**Fig. 5.**
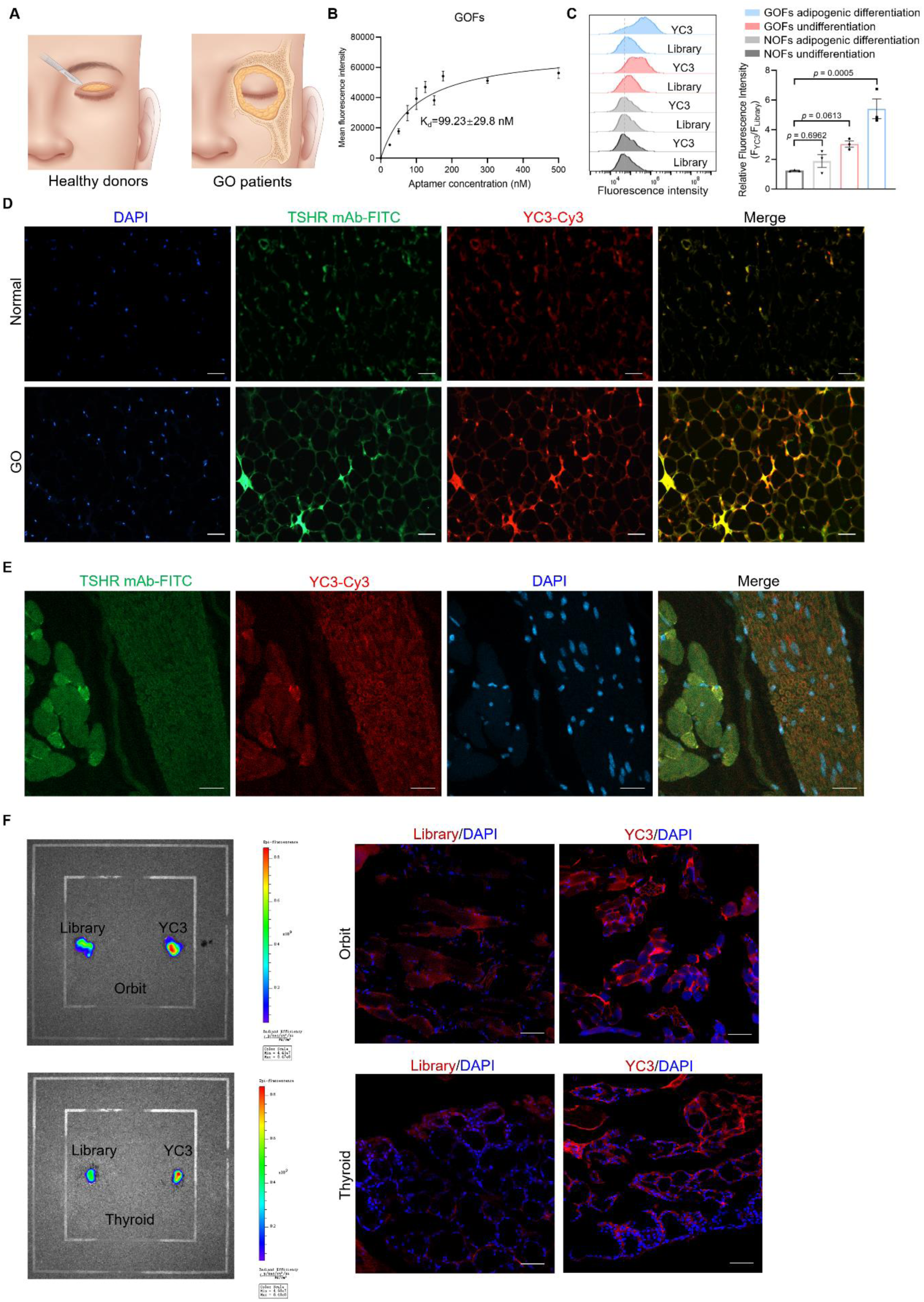
Aptamer YC3 can target the orbit. (**A**) Schematic illustration of orbital adipose tissues from GO patients who underwent orbital decompression surgery, and healthy donors who underwent blepharoplasty. (**B**) The dissociation constant of YC3 for Graves orbital fibroblasts (GOFs) was determined by flow cytometry, using the following equation: Y=B_max_ X/(K_d_+X) (X representing the concentration of aptamer, Y representing the mean fluorescence intensity, and B _max_ representing the peak fluorescence intensity), n=3 independent samples. (**C**) Flow cytometry analysis of the binding of YC3 to Graves orbital fibroblasts (GOFs) and normal orbital fibroblasts (NOFs) with or without adipogenic differentiation. Quantitative analysis of the relative fluorescence intensity, n=3 independent samples in each group. Data are represented as mean±SEM. One-way ANOVA, followed by Tukey’s multiple post hoc test was used to calculate *P* values. (**D**) Confocal microscopy was used to analyze the colocalization of DAPI stain, anti-TSHR antibody and YC3 on orbital adipose tissue sections from healthy donors and patients with GO. Representative images were shown. Scale bars: 50 μm. n=6 independent samples. (**E**) Confocal microscopy was used to analyze the colocalization of DAPI stain, anti-TSHR antibody and YC3 on orbital adipose tissue sections from BALB/c mice. Representative images were shown. Scale bars: 20 μm. n=3 independent samples. (**F**) Fluorescent imaging of orbits dissected from BALB/c mice after intravenous injection of Cy5-labeled YC3 or Library, and frozen section fluorescence imaging of Cy5-labeled aptamers in the corresponding orbits and thyroids. The sections were counterstained with DAPI. Representative images were shown. Scale bars: 20 μm.

To determine whether the specific binding of YC3 to TSHR can be as a molecular probe for clinical application, orbital adipose tissues from GO patients and healthy donors were incubated with Cy3-labeled YC3 and FITC-labeled TSHR antibodies. Compared to healthy donors, orbital adipose tissues from GO patients showed a higher expression of TSHR, as well as a stronger binding of YC3 (Fig. 5D). Meanwhile, YC3 and TSHR showed strong colocalization in orbital adipose tissues. Moreover, we found that TSHR was expressed in mouse orbital tissues, with TSHR expression in the thyroid serving as a positive control (Fig. S7A-B). Strong colocalization of YC3 and TSHR was observed in the orbital tissues of mice (Fig. 5E). To investigate the targeting ability of YC3 *in vivo*, we intravenously injected an equal amount of Cy5-labeled YC3 or Cy5-labeled Library via the tail vein. Subsequent fluorescent imaging of dissected mouse orbits and thyroids (serving as positive controls) and corresponding frozen tissue sections revealed that the aptamer YC3 exhibited a stronger fluorescence signal in both orbital and thyroid tissues compared to the aptamer Library (Fig. 5F). Taken together, YC3 demonstrates great targeting performance in both human and mouse orbital tissues, indicating its promising potential for *in vivo* application as a molecular probe.

### Subhead 6: Aptamer YC3 inhibits M22-stimulated activation of orbital fibroblasts

Activated orbital fibroblasts exhibit enhanced contractile and secretory activities^26,27^. In GO, this enhancement specifically promotes inflammation cytokines and extracellular matrix (ECM) production, leading to persistent orbital inflammation, fibrosis and pathological tissue remodeling^28–32^. Both TSH and M22 significantly increased IL-6 and IL-8 mRNA and protein levels in GOFs. M22, but not TSH, also elevated IL-1β mRNA and protein levels (Fig. S6A-B). Hyaluronic acid (HA) is a key component of the ECM, and its dysregulation is implicated in inflammatory and fibrotic diseases^33^. Therefore, we also examined the effects of TSH and M22 on hyaluronic acid synthetases (HAS1, HAS2 and HAS3) in GOFs. The results showed that M22, but not TSH, significantly increased the mRNA levels of these synthetases and HA levels (Fig. S6C-D).

To further investigate the functional effects of YC3 on GOFs, we examined several key cellular processes. Fibroblast activation promotes tissue remodeling in fibrotic pathologies through enhanced contractile force and migratory activity. Actin-myosin interactions, particularly stress fibers, are central to generating mechanical tension in fibrosis^34^. Our results demonstrated that YC3 mitigated an M22-induced increase in actin fiber density and prominent stress fiber formation in GOFs (Fig. 6A). Nuclear translocation of NF-κB p65 plays a key role in the inflammatory response by promoting the production of inflammatory cytokines and chemokines. YC3 abrogated M22-induced nuclear translocation of NF-κB p65 (Fig. 6B). We also investigated the effects of YC3 on the production of inflammatory cytokines and HA. As expected, YC3 significantly reduced the mRNA and protein levels of IL-6, IL-8, and IL-1β in M22-stimulated GOFs. Furthermore, YC3 inhibited HAS mRNA expression in GOFs and reduced HA levels in the GOFs supernatant. The anti-inflammatory and anti-ECM production effects of YC3 were comparable to those of the clinical drug Teprotumumab (Fig. 6C-D).

**Fig. 6.**
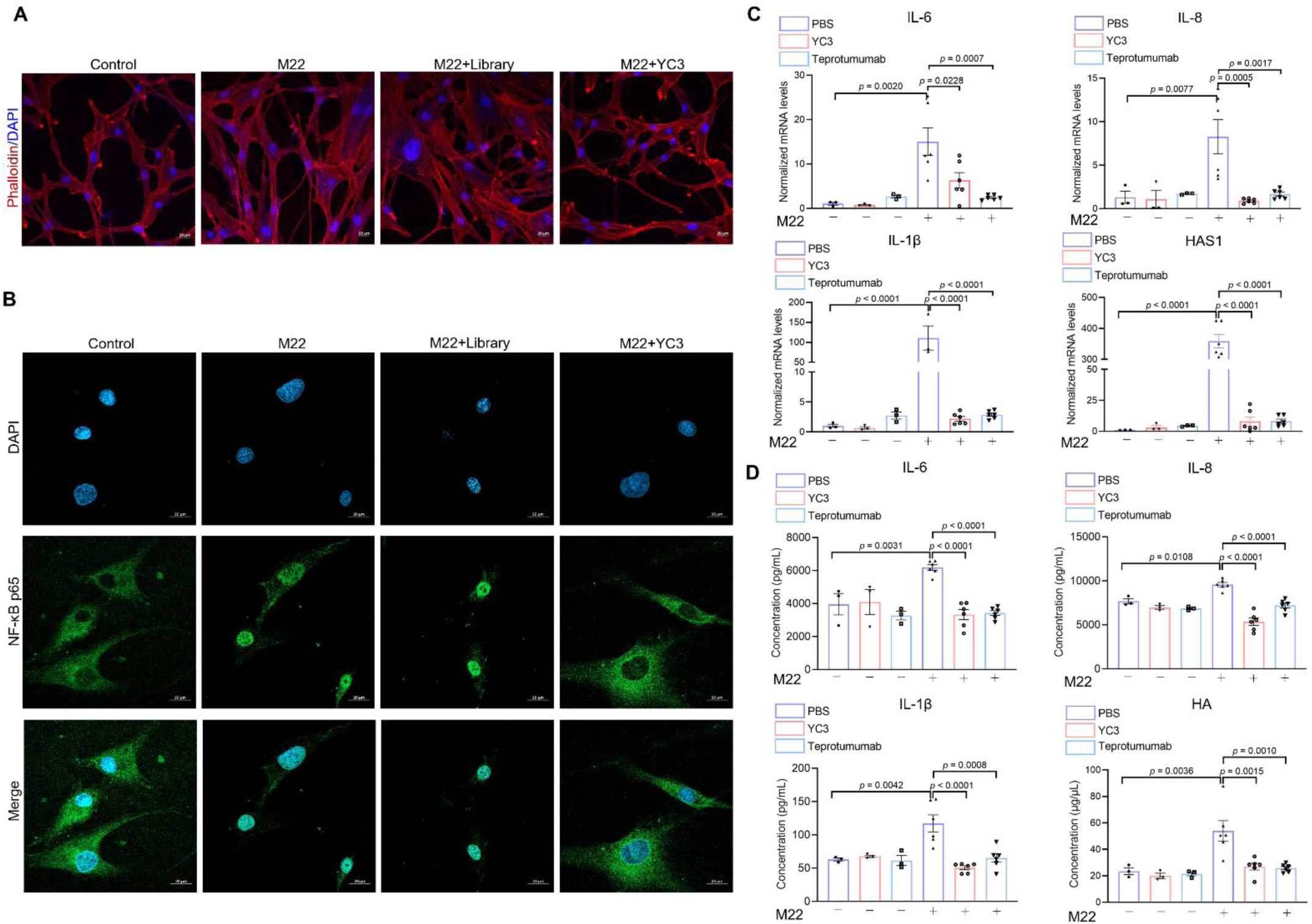
Aptamer YC3 can inhibit the activation of orbital fibroblasts stimulated by M22. (**A**) Phalloidin staining for actin filaments in GOFs stimulated by M22 and treated with aptamers Library or YC3 (2 μM). The cells were counterstained with DAPI. Representative images were shown. Scale bars: 20 μm. The experiments were repeated three times independently. (**B**) Fluorescence image of NF-κB p65 in GOFs stimulated by M22 and treated with aptamers Library or YC3 (2 μM). Representative images were shown. Scale bars: 20 μm. The experiments were repeated three times independently. (**C**) The mRNA expression levels of IL-6, IL-8, IL-1β, and HAS1 in GOFs stimulated by M22 and treated with aptamer YC3 (2 μM) or Teprotumuab (5 μg/mL) by qPCR, n=3 independent samples in non-stimulated group, n=6 independent samples in M22-stimulated group. Data are represented as mean±SEM. One-way ANOVA, followed by Tukey’s multiple post hoc test was used to calculate *P* values. (**D**) The concentrations of IL-6, IL-8, IL-1β, and HA in the supernatant medium of GOFs stimulated by M22 and treated with aptamer YC3 (2 μM) or teprotumumab (5 μg/mL) were measured by ELISA, n=3 independent samples in non-stimulated group, n=6 independent samples in M22-stimulated group. Data are represented as mean±SEM. One-way ANOVA, followed by Tukey’s multiple post hoc test was used to calculate *P* values.

### Subhead 7: Aptamer YC3 improves outcomes in a GO mouse model

To investigate the role of YC3 *in vivo*, we established a GO mouse model by intramuscular injection of adenovirus expressing the TSHR α-subunit (Ad-TSHR289), as previously reported^35^. During GO model construction, the body weight of mice in both the MOCK and GO groups increased over time. However, the GO group gained more weight compared to the MOCK group (Fig. S8A). At week 18, the serum concentrations of T3, T4, and thyrotropin receptor antibody (TRAb) in the GO group were significantly higher than those in the MOCK group (Fig. S8B). For ocular symptoms, mice in the GO group exhibited signs of inflammation, including eyelid broadening, eyelid swelling, and congestion, which resembled the symptoms observed in GO patients (Fig. S8C). To monitor inflammatory events and remodeling of orbital tissues *in vivo*, we performed serial non-invasive Magnetic Resonance Imaging (MRI) on living mice. T2-weighted MRI was used to image distinct orbital tissue types, including extraocular muscle, adipose tissue, optic nerve, and all anatomical structures. As shown in Figure S8D, the MRI images from the GO group revealed extraocular muscle hypertrophy and high signal intensity in the retrobulbar area, indicating inflammation and remodeling of the orbital tissues. These findings are similar to the imaging changes observed in GO patients.

As shown in the timeline (Fig. 7A), after seven injections of the adenovirus, mice with GO were randomly divided into four groups: periorbital injection with PBS (GO+Blank group); periorbital injection with the modified aptamer Library (PS-Library-PEG, referred to as Apt-Library) (GO+Apt-Library group); periorbital injection with the modified aptamer YC3 (PS-YC3-PEG, referred to as Apt-YC3) (GO+Apt-YC3 group); periorbital injection with dexamethasone sodium phosphate (DSP) (GO+DSP group), serving as a positive control. All injections were administered three times a week. During the treatment, the weight of mice remained stable, except for those in the GO+DSP group, which gained weight during the treatment (Fig. 7B). At week 24, the end of treatment, the concentrations of T3, T4, and TRAb in the serum of mice from the GO+Apt-YC3 group were significantly reduced compared with the GO+Apt-Library group. In T2-weighted MRI images, compared with the GO+Apt-YC3 group, extraocular muscle hypertrophy and orbital tissue inflammation and swelling were more severe in the GO+Blank group and the GO+Apt-Library group (Fig. 7D), indicating that YC3 improved the pathological remodeling of orbital tissues.

**Fig. 7.**
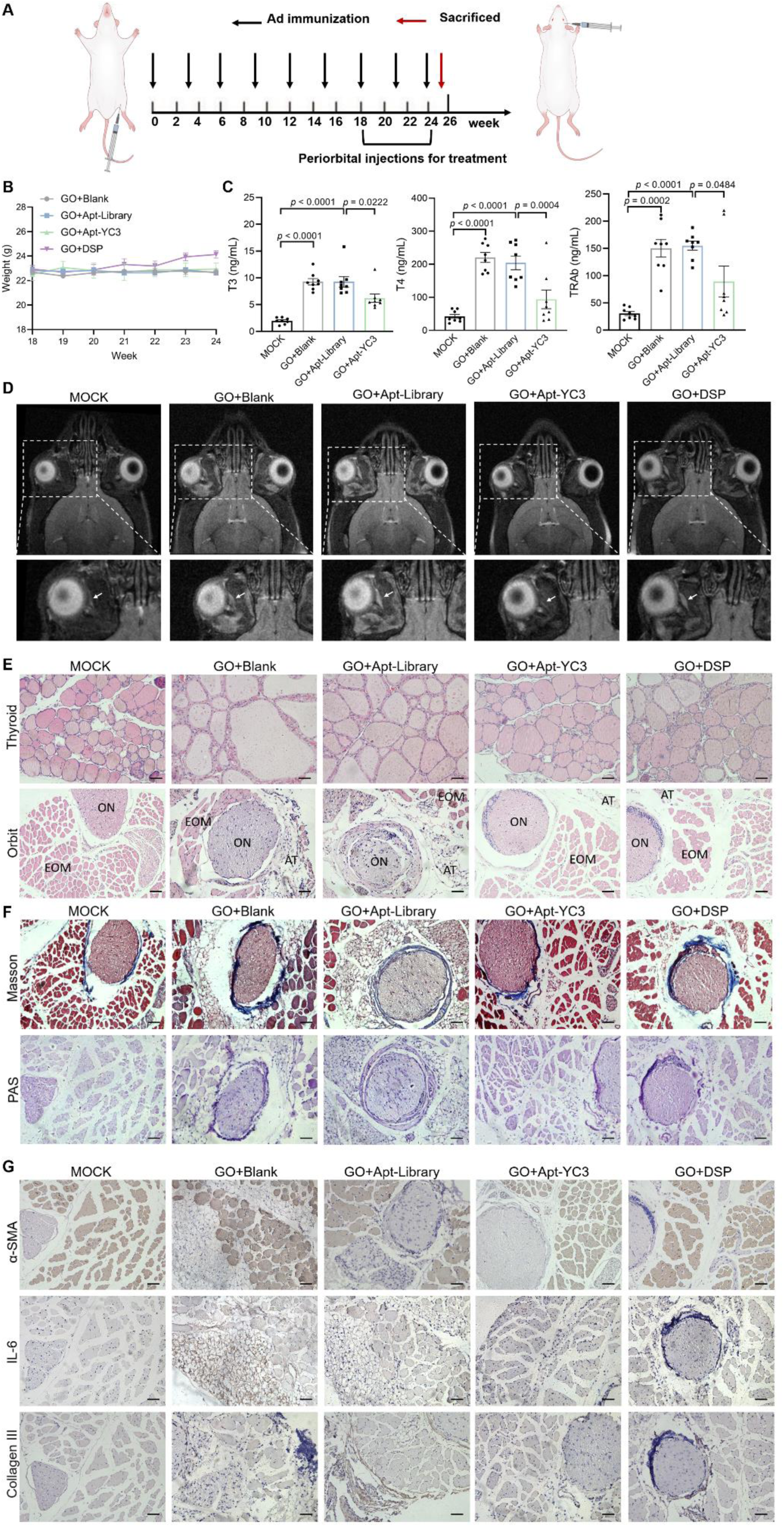
Aptamer YC3 can improve the outcome of GO mouse model. (**A**) Experimental timeline for model construction and treatment of mice. Mice were injected with Ad-TSHR289 or Ad-MOCK, and then mice injected with Ad-TSHR289 were randomly grouped, followed by treatment with Apt -YC3, Apt-Library, or DSP, respectively. (**B**) Measurement of the weight of mice in four groups once a week following the treatment. Data are represented as mean±SEM. (**C**) The concentration of T3, T4, and TRAb in mice serum in four groups by ELSIA, n=8 independent samples in each group. Data are represented as mean±SEM. One-way ANOVA, followed by Tukey’s multiple post hoc test was used to calculate *P* values. (**D**) Representative images of T2-weighted Magnetic Resonance Imaging of the mice from five groups. The lower images provide an enlarged view of the areas within the white dashed boxes shown in the corresponding upper images. White arrows denote the extraocular muscle. (**E**) Representative images of Hematoxylin-eosin (H&E) staining from thyroid and orbital tissue sections for the MOCK group, the GO+Blank group, the GO+Apt-Library group, the GO+Apt-YC3 group, and the GO+DSP group. ON, optic nerve; EOM, extraocular muscle; AT, adipose tissue. Scale bars: 20 μm. (**F**) Representative images of Masson staining and periodic acid-schiff (PAS) staining from orbital tissue sections for the MOCK group, the GO+Blank group, the GO+Apt-Library group, the GO+ Apt-YC3 group, and the GO+DSP group. Scale bars: 20 μm. (**G**) Representative images of α-SMA, IL-6, and Collagen Ⅲ staining from orbital tissue sections for the MOCK group, the GO+Blank group, the GO+Apt-Library group, the GO+ Apt-YC3 group, and the GO+DSP group. Scale bars: 20 μm.

Histopathological analysis was employed to assess the thyroid and orbital histological features caused by adenovirus TSHR289 immunization and the effect of YC3 on TSHR-immunized mice. As shown in Figure 7E, thyroid sections from the GO+Blank group revealed diffuse enlargement of the thyroid gland. The thyroid follicles exhibited cuboidal to cylindrical follicular cells with a small amount of colloid, indicating hyperactivity. Orbital sections from the GO+Blank group showed inflammatory cell infiltration, expanded orbital adipose tissue, and disordered muscle fibers. Interestingly, in the GO+Apt-YC3 and GO+DSP groups, YC3 and DSP alleviated the morphological and histological changes in the orbit, reduced inflammatory cell infiltration, decreased adipose tissue expansion, and promoted the alignment of muscle fibers. Fibrosis is characterized by excessive and abnormal accumulation of extracellular matrix, which can impair tissue function^36^. The degree of fibrosis in orbital tissues was determined by the concentration of retrobulbar collagen, acidic mucin, and glycosaminoglycans^37^. Normal mice had very little collagen and glycogen in their orbital tissues. However, in the GO+Blank group and the GO+Library group, collagen and glycogen deposition increased significantly around the optic nerve and extraocular muscle bundles. Notably, YC3 reversed the accumulation of collagen and glycogen, thereby mitigating the fibrotic response in the orbit (Fig. 7F, S10A-B). We further investigated the levels of fibrosis markers such as alpha-smooth muscle actin (α-SMA) and collagen III and the inflammatory cytokine IL-6 in the orbital tissues of mice. Our findings demonstrated that in the GO+Blank group and the GO+Library group, there was an increase in the levels of α-SMA, IL-6, and collagen III. Conversely, YC3 significantly reduced these expressions (Fig. 7G, S10C-E). Taken together, these results demonstrate that aptamer YC3 significantly improves the outcomes of GO mice by alleviating hyperthyroidism, reducing inflammation, decreasing ECM accumulation, and mitigating tissue fibrosis.

## Discussion

Clinical studies have confirmed the pathological role of TSHR in GO ^10, 38, 39^, but the therapeutic efficacy of TSHR inhibition for GO remains uncertain. Given the critical role of TSHR in metabolism, knocking out TSHR would cause significant physiological impairments ^40^. Therefore, studies of TSHR intervention in GO primarily adopt pharmacological approaches. A clinical trial reports that TSHR monoclonal antibodies, K1 -70, effectively improve symptoms of GO in one case^41^. The limited and preliminary *in vivo* observations suggest that TSHR inhibition may hold promise for the treatment of GO. In this study, we obtained TSHR-targeting aptamer YC3 with nanomolar affinity and inhibitory potency. YC3 can serve as a molecular probe for TSHR, effectively detecting TSHR expression in cell lines and living tissues. In GOFs, YC3 reduced TSAbs-induced activation, decreasing inflammatory factor production and ECM secretion. Its *in vitro* efficacy was comparable to that of Teprotumumab, an anti-IGF1R monoclonal antibody, which has demonstrated significant efficacy in clinical trials^42–44^. It would be interesting to further explore the efficacy of YC3 and Teprotumumab combination as a multi-target therapy for GO. In a GO mouse model, YC3 reduced the levels of TRAb, potentially by regulating the immune system and restoring immune tolerance. We found that YC3 not only alleviates the pathological remodeling of the orbit but also relieves hyperthyroidism. This dual benefit highlights YC3’s potential in treating both the ocular and systemic manifestations of Graves’ disease. Collectively, this study provides the first evidence that pharmacological inhibition of TSHR has a therapeutic effect in GO mouse models and offers a promising therapeutic molecular tool for GO and other TSHR-related diseases.

Among the reported TSHR inhibitors, both orthosteric and allosteric inhibitors have been identified. The orthosteric inhibitor K1 -70 targets the same binding sites on the TSHR as TSH and TSAbs ^45^. Allosteric inhibitors include some small molecule compounds, but research on the binding sites and mechanisms of these allosteric inhibitors is limited. For instance, Neumann and Latif have investigated the allosteric inhibitors Org41841 and VA-K-14 using molecular docking approaches ^46, 47^. The results indicate that allosteric inhibitors may target a pocket formed by the seven transmembrane helices and three extracellular loops of the TSHR, maintaining the receptor in an inactive conformation^4^. These allosteric inhibitors block signal transduction from the extracellular to the transmembrane region, thereby preventing the conformational changes required for TSHR activation. However, the potential allosteric inhibitory mechanisms identified so far lack experimental validation, and it is unclear whether other allosteric sites can influence TSHR activation. In this study, we delved into the molecular mechanism of the interaction between the aptamer YC3 and TSHR. Aptamers are single-stranded nucleic acids with flexible structures, making it challenging to characterize their interaction sites. Surprisingly, in our work, we attempted to identify the binding sites of YC3 to TSHR through competition experiments, computational structural analysis, and verification through site-directed mutagenesis studies. Interestingly, YC3 binding does not compete with TSAbs. While TSAbs bind to the concave face of the leucine-rich repeat region in the extracellular domain, YC3 binds to the convex face of the same region. We identified K218, D219, K244, and N170 as key amino acid residues contributing significantly to YC3’s binding energy, with K218 being the most critical. This suggests that YC3 acts as an allosteric inhibitor. To our knowledge, allosteric modulators of TSHR targeting these sites have not been previously reported. We speculate that the binding of YC3 to TSHR alters the receptor conformation, making it unfavorable for signal transduction. Future studies could use cryo - electron microscopy to investigate YC3’s impact on TSHR conformation. While K218 significantly influences YC3 binding, its role in TSHR activation remains unclear and warrants further investigation as a potential target for allosteric inhibitors. We found that the YC3 -TSHR interaction involves an extensive interface, with a quadruple mutant only partially affecting YC3 binding. This large contact area includes many functional groups, resulting in strong binding forces and high specificity. Our findings suggest a complex regulatory mechanism that merits further exploration. Overall, our study provides a molecular foundation for YC3’s binding and inhibitory functions, offering the newly identified allosteric site for developing effective TSHR inhibitors and potentially transforming therapeutic strategies for TSHR-related disorders.

Aptamer inhibitors have gained significant attention and spurred active development due to their advantages and clinical success. Most aptamers function by competitively binding to target receptors, blocking ligand - receptor interactions and modulating cellular signaling ^22, 48–50^. This competitive mechanism can limit efficacy, particularly at high concentrations of endogenous ligands, and often leads to poor selectivity. For example, inhibitors targeting ATP-binding sites can affect multiple kinases, resulting in off-target effects. Consequently, these inhibitors may cause significant adverse side effects ^51^. Allosteric modulators bind to sites distinct from orthosteric sites, offering higher specificity and selectivity for specific protein subtypes. This reduces side effects and enables effectiveness at lower doses. As a result, allosteric modulators present a promising avenue for developing novel therapeutic agents. YC3, as a unique allosteric inhibitor, combines the advantages of allosteric modulators and serves as a new paradigm for the development of functional aptamers. However, aptamers’ single-stranded nucleic acid structure makes them susceptible to nuclease degradation and rapid *in vivo* elimination, complicating treatment for chronic conditions like GO, where frequent administration is impractical. In this work, simple chemical modifications to YC3 improved its *in vivo* stability while maintaining its binding affinity, highlighting its potential for translation into a therapeutic agent. Fo r future clinical applications, strategies like polyvalent aptamers, aptamer conjugates, and aptamer hydrogels hold promise for improving drug loading and extending the half-life of therapeutic agents^52^. To advance YC3 for clinical use, further modifications and optimizations are needed, along with thorough toxicity studies, to enhance its stability, efficacy, and *in vivo* safety.

In summary, this study developed YC3, a nucleic acid aptamer with high affinity and specificity for TSHR. YC3 shows robust inhibitory pharmacodynamic activity both *in vitro* and *in vivo*. Unlike endogenous ligands, YC3 binds to key residues (K218, D219, K244, and N170) on the convex surface of the LRD in the extracellular region of TSHR, inhibiting receptor activation through allosteric effects. This reveals a unique target site for designing selective allosteric inhibitors. Additionally, YC3 effectively alleviates GO symptoms in a mouse model, providing the first preclinical evidence that pharmacological inhibition of TSHR has a therapeutic effect in GO. Our work underscores YC3’s potential as a therapeutic agent for GO and other TSHR-related diseases, offering promising precision therapy.

## Materials and Methos

### Materials

Yeast tRNA was purchased from Sigma-Aldrich (Shanghai, China). Recombinant human TSH alpha/beta heterodimer protein was purchased from the R&D System (Minneapolis, USA). Human monoclonal autoantibody to the TSHR (M22) was purchased from RSR (Cardiff, UK). Teprotumumab was purchased from AntibodySystem (Strasbourg, France). All other reagents were purchased from Beyotime (Shanghai, China) unless stated otherwise.

### Ethical statement

Human tissue specimens were obtained with approval from the Eye Hospital of Wenzhou Medical University Ethics Committee (Approval No. 2023-074-K-62), and signed informed consent was obtained from all participants. All animal experiments were conducted in accordance with the guidelines of the Eye Hospital of Wenzhou Medical University and approved by the Institutional Animal Care and Use Committee (Approval No. YSG23110303).

### Human orbital fibroblast isolation and culture

Orbital adipose tissues were obtained from six patients with GO who underwent orbital decompression surgery for severe GO, and six healthy donors who underwent blepharoplasty. None of the GO patients had received steroid therapy or radiotherapy for at least three months prior to surgery. All surgeries were performed at the the Eye Hospital of Wenzhou Medical University, Wenzhou, China. Patient details and histories of antithyroid medication use are provided in Table S3.

Orbital fibroblasts (OFs) were isolated by mincing tissue pieces and plating them in culture dishes, allowing time for adhesion before immersing in high-glucose DMEM with 20% FBS and 1% penicillin/streptomycin (P/S). Experiments were conducted with strains between passages 3 and 7, repeated with at least three independent strains.

### Cell culture

All the cell lines and cell culture reagents are listed and summarized in Table S4. All cell lines were cultured at 37°C in a humid atmosphere with 5% CO_2_.

### Adipogenesis

When the OFs were confluent to 100%, the complete growth medium was converted to a differential medium (DM). The base medium for DM was high-glucose DMEM containing 10% FBS and 1% P/S, supplemented with 0.1 mM indomethacin (Sigma), 0.1 μM dexamethasone (Sigma), 0.5 mM 3 -isobutyl-1-methylxanthie (IBMX) (MedChemExpress), and 10 μg/mL insulin (Procell). The medium was refreshed every 4 to 5 days, and the induction period lasted for 10 -14 days, as previously described ^53^.

### Expression vector construction and transfection

Human TSHR cDNA (GeneCopoeia, NM_000369.4) was cloned into a pLVX-Puro vector (oe-TSHR) and verified by DNA sequencing. The wild-type TSHR plasmid was co-transfected with lentiviral packaging vectors to produce recombinant lentiviral particles, which were used to infect HEK293T cells, followed by puromycin selection to establish TSHR - overexpressing HEK293T cells. An empty plasmid was similarly packaged into lentiviral particles to generate MOCK cells. Mutant TSHR plasmids were constructed by RIBOBIO CO., Ltd. and transfected into HEK293T cells using Lipofectamine™ 3000 (Invitrogen, USA).

### DNA Library, SELEX primers, and buffers

The ssDNA Library used by Cell-SELEX contained a 42-nucleotide central random region flanked by two 19 -nucleotide primer-binding regions. A FAM-labeled primer (5’-FAM-ACCGACCGTGCTGGACTCA-3’) and a biotin-labeled primer (5’-biotin-CGCCAGGCTCGCTCATAGT-3’) were utilized for PCR amplification. The washing buffer was composed of DPBS containing 5 mM of MgCl_2_ and 4.5 g/L of glucose. The binding buffer was formed from the washing buffer containing 0.1 mg/mL yeast tRNA and 1 mg/mL BSA. All DNA sequences listed in Table S1 were HPLC-purified and synthesized in Sangon Biotech (Shanghai, China).

### Cell-SELEX Procedure

Cell-SELEX procedures followed published protocols ^54^. Initially, 10 OD of the ssDNA library was dissolved in binding buffer, denatured at 95°C for 10 minutes, and cooled on ice. In the first round, ssDNA was incubated with TSHR-293T cells at 4°C for 90 minutes, washed three times, and the bound ssDNA was released by heating at 95°C for 10 minutes. The harvested ssDNA was amplified via PCR using FAM-labeled forward and biotin-labeled reverse primers, captured using streptavidin-coated sepharose beads (GE Healthcare) and isolated after denaturation. From the third round, negative selection with MOCK cells was introduced to enhance specificity. The ssDNA pool was incubated with MOCK cells for 30 minutes, and unbound ssDNA was collected for positive selection. As the screening rounds progressed, the criteria became increasingly stringent to enhance the affinity and specificity of the sequences. After enrichment, high -throughput sequencing was performed using Illumina MiSeq by Sangon Biotech (Shanghai, China).

### Flow cytometric analysis

Flow cytometry analysis was used to assess the binding ability of evolved ssDNA pools, aptamers, or antibodies to cells. TSHR -293T and MOCK cells were incubated with FAM-labeled ssDNA pools at 4°C for 30 minutes, washed, and resuspended in the washing buffer for flow analysis. The K _d_ of the aptamer was calculated using the following equation: Y=B _max_ X/(K_d_+X) by GraphPad Prism 8.0. TSHR expression on various cell membranes was evaluated by incubating cells with FITC-labeled IgG2a, followed by an anti-TSHR antibody, and FITC-conjugated secondary antibodies Cell fluorescence signals were recorded using a FACS Calibur flow cytometer (BD Bioscience) and analyzed with FlowJo 10.0. All experiments were repeated three times.

### Immunofluorescence analysis assay

Immunofluorescence analysis mainly included the binding ability of aptamers or antibodies to cells or tissues. For aptamers, cells were incubated with fluorescence-labeled aptamer in 500 µL of binding buffer at 4°C for 30 minutes. For antibodies, cells were fixed with 4% paraformaldehyde, washed, and blocked with 10% normal goat serum. Cells were then incubated with primary antibodies overnight at 4°C. After washing, secondary fluorescent antibodies were incubated, followed by DAPI nuclear counterstaining. Images were acquired using a Cell Observer SD system (Zeiss, Germany).

Paraffin-embedded tissue sections were prepared, heated at 60°C, deparaffinized in xylene, and rehydrated through decreasing ethanol concentrations. Epitope retrieval was performed using citrate buffer (pH 6.0). Sections were blocked with DNA blocking buffer (20% FBS, 0.1 mg/mL salmon sperm DNA), then incubated with anti-TSHR antibody (1:500, Abcam, ab27974) overnight at 4°C. After washing, sections were stained with fluorescence-conjugated secondary antibodies. Sections were then incubated with Cy3-labeled aptamer Library or YC3, counterstained with DAPI, sealed, and imaged using a fluorescence microscope (Leica, Bannockburn, IL, USA).

### Aptamer-mediated pull-down assay

The pull-down assay was conducted as previously described ^55^. 3×10^7^ TSHR-293T cells were dissociated with 0.2% EDTA, collected by centrifugation, and membrane proteins were extracted using a kit (PH0710, Phygene). The membrane proteins were divided into four parts. Three of these parts were incubated in a DNA-blocking buffer at 4°C. Two of these parts were then incubated with biotin-labeled aptamer Library (control) or biotin-labeled aptamer YC3 overnight at 4°C. Then these three parts were incubated with streptavidin-coated magnetic beads at 4°C. After centrifugation and washing, the samples were treated with protein loading buffer, heated, and subjected to SDS-PAGE electrophoresis followed by Western blotting.

### Small interfering RNA interference and transfection

Small interfering RNA (siRNA) interference was conducted according to the published protocol^56^. siRNA sequences targeting TSHR and a random sequence (listed in Table S5) were synthesized by RIBOBIO CO., Ltd. (Guangzhou, China). In brief, TSHR-293T cells were transfected with siRNA by Lipofectamine™ 3000 transfection reagent (Invitrogen, USA).

### Measurement of cAMP production

Cells were seeded in 24-well plates and starved with 1% FBS overnight. The next day, cells were incubated in antibiotic-free serum-free media containing 0.5 mM IBMX to avoid cAMP decay, followed by adding aptamers, with or without M22 (100 ng/mL). The intracellular extracts were gathered and stored at -80°C until the cAMP analysis was conducted using the cAMP assay (KGE002B, R&D Systems, USA). The assay employs polyclonal antibodies to competitively bind cAMP in both standard and sample extracts, as described previously ^57^.

### RNA Isolation, reverse transcription and quantitative RT-PCR

Total RNA from cells and adipose tissues was extracted using TRIzol reagent (Invitrogen) following the manufacturer’s instructions. RNA concentration was measured with a UV spectrophotometer. cDNA was synthesized using a reverse transcription kit (Promega, USA), and real -time PCR was conducted with PCR SYBR Green (APExBIO, USA). Three independent experiments were performed to ensure reliability. Primer pairs were synthesized by Tsingke Biotech and detailed in Supplementary Table S6.

### Western blotting

Total protein was extracted using RIPA buffer (EpiZyme) with protease inhibitor cocktail, and concentrations were determined using a protein assay reagent (EpiZyme). Protein samples were separated on SDS -PAGE gels and transferred to nitrocellulose membranes using a Bio-Rad Western blot transfer system. Membranes were washed with TBST, blocked with 5% skim milk and incubated with primary antibodies overnight at 4°C. After washing, membranes were incubated with Goat anti-Mouse IgG (1:5000, LI-COR, IRDye680CW). Detection was performed using an Odyssey infrared scanning system (LI-COR). Primary antibodies used included: anti-TSHR (Abcam), anti-β-actin (Proteintech), anti-GAPDH (Bioss), anti-ERK1/2 (Cell Signaling Technology), anti-phospho-ERK1/2 (Cell Signaling Technology), anti-MEK1/2 (Cell Signaling Technology), anti-phospho-MEK1/2 (Cell Signaling Technology), anti-AKT (Proteintech), and anti-phospho-AKT (Cell Signaling Technology). All primary antibodies were used at a dilution of 1:1000.

### Enzyme-linked immunosorbent assay

The concentrations of inflammatory cytokines (IL-6, IL-8, and IL-1β) in cell-conditioned supernatants were measured using ELISA kits (FineTest, Wuhan, China): EH0201 for IL-6, EH0205 for IL-8, and EH0185 for IL-1β, following the manufacturer’s protocols. Similarly, T3, T4, and TRAb concentrations in mouse serum from the orbital venous sinus were measured using ELISA kits (FineTest): EU0403 for T3, EU0402 for T4, and EM1423 for TRAb. The concentration of HA was quantified using a specific ELISA kit from Echelon Biosciences (Salt Lake City, UT), according to the manufacturer’s instructions.

### Surface plasmon resonance experiments

Surface Plasmon Resonance (SPR) was used to determine the binding affinity between TSHR and the aptamer YC3 using a BIAcore 8K instrument with CM5 chips at 25°C. Anti-His antibodies (Proteintech, 66005 -1-lg) were immobilized on the CM5 chip via amine coupling to create an anti -His CM5 chip. TSHR-His recombinant proteins (Sangon Biotech, China) were then captured by this chip. YC3 was serially diluted to concentrations ranging from 0 to 200 nM and injected at 30 μL/min for 180 seconds, followed by 180 seconds of dissociation. All experiments were conducted in running buffer (DPBS with 5 mM Mg²⁺, pH 7.4). The relative response was recorded and fitted to a 1:1 kinetic model using Biacore evaluation software.

### PEGylation of aptamer

PEGylation was performed according to previously described methods ^58^. Specifically, the thiol-labeled aptamer was suspended in PBS and reduced with 20 mM DTT (PHYGENE, China). After the reaction, the DTT was removed using spin filters. The reduced aptamers were then incubated with an excess amount of 10 kDa maleimide-terminated polyethylene glycol (Mal-PEG) (Ponsure Biotechnology, Shanghai, China). Following conjugation, unconjugated Mal-PEG was removed using spin filters.

### Structures of Protein and Aptamer

The protein structures of TSHR and M22 were obtained from the RCSB Protein Data Bank (PDB IDs: 7XW5, 7XW6) ^2^. The Vienna number of aptamer YC3 was generated online using the MFOLD web server ^59^, and the corresponding three-dimensional structure with the best-predicted energy was constructed using 3dRNA/DNA ^60^. To eliminate steric conflicts within the aptamer system, a position-constrained molecular dynamics simulation was performed.

### Molecular docking

Aptamer YC3 and protein TSHR were subjected to molecular docking using Rosetta software^61^, involving two steps: aggressive sampling with a centroid model followed by fine optimization using a full -atom model. Initially, rigid-body blind docking generated at least 2,000 conformations with position-restrained heavy atoms and randomly oriented protein and aptamer. Amino acids forming frequent hydrogen bonds (>0.4 frequency) were selected as potential binding sites, excluding distant sites. In the fine optimization step, the aptamer was positioned within 10 Å of the binding pockets and perturbed by a 3 Å translation and an 8°rotation before each simulation. Side chains of protein residues and the complete DNA were allowed to move. Up to 100 conformers were assessed, and the one with the lowest binding energy was selected for subsequent molecular dynamics simulation.

### Molecular dynamics simulations

The optimal docking conformation of the TSHR-YC3 complex from Rosetta docking served as the starting point for molecular dynamics (MD) simulations using Amber 20 software ^62^. Simulations lasted at least 100 ns, with RMSD monitored until system equilibrium was achieved. The protein system utilized the GAFF and ff14SB force fields, enclosed in a cubic water box with a 10 Å radius around the protein, and Na+ ions were added to neutralize the system. The simulation process included: (1) Two -step energy minimization; (2) System equilibration; and (3) Dynamics simulation. The cutoff distance for van der Waals and short -range electrostatic interactions was set to 10 Å, and the Particle -Mesh-Ewald method was used to calculate long-range electrostatic interactions. In this system, the average structure from the time interval of 60 -80 ns during the equilibrium state was extracted for subsequent binding mode analysis.

### Establishment of GO mouse model by immunization with Ad -TSHR289

Female BALB/c mice were purchased from Weitong Lihua Experimental Animal Technology Co., Ltd. and housed in a pathogen-free environment at the Eye Hospital of Wenzhou Medical University. Mice were randomly divided into three groups: GO, MOCK, and Blank. The GO group received intramuscular injections of 2 × 10^9^ particles of Ad-TSHR289 in 50 μL PBS, the MOCK group received an equal number of particles of empty -vector adenovirus in 50 μL PBS, and the Blank group received 50 μL PBS. Injections were administered every three weeks for a total of nine injections, with all mice sacrificed at week 25. The mice were maintained in a controlled environment with a 12 -hour light/dark cycle, had unrestricted access to water and food, and were weighed every three weeks.

### Administration regimen in GO mice

After constructing the GO models for 16 weeks, T3, T4, and TRAb serum levels in the GO group were measured, with successful model construction defined as levels more than twice those of the Blank group. Successfully modeled GO mice were randomly divided into four groups. From weeks 18 to 24, mice were anesthetized with isoflurane and received periorbital injections three times per week. Each eye was injected with either PBS, aptamer PS-YC3-PEG (10 mg/kg), aptamer PS-Library-PEG (10 mg/kg), or DSP at 1.5 mg/kg. The treatment regimen is illustrated in Fig. 7A. The appearance of the mice’s eyes was observed and photographed, and their body weight was recorded weekly during the treatment period.

### Magnetic Resonance Imaging

MRI was performed using a 9.4 Tesla animal scanner (BioSpec 94/20 USR, Bruker BioSpin, Ettlingen, Germany) with ParaVision 3.2 software. A 3 -channel phased-array surface coil (Bruker) served as the receiver, and an 86-mm diameter volume coil (Bruker) was used as the transmitter. T2 -weighted images were acquired using a Turbo RARE sequence with the following parameters: Field of View=2.0 × 2.0 cm², Matrix Size=256 × 256, Slice Thickness=0.30 mm, Interslice Distance=0.0 mm, Number of Slices=20, TR/TE=2500/33 ms.

### Histopathological examination

For H&E staining, sections were stained with hematoxylin, rinsed, differentiated, and rinsed again. Slides were then immersed in ammonia solution, rinsed, dehydrated with 85% and 95% ethanol, and stained with eosin. After dehydration with a gradient of alcohol, slides were cleared with xylene and sealed with neutral gum. For Masson staining, follow the instructions provided with the Masson staining kit (Beyotime, China). For periodic acid-schiff (PAS) staining, follow the instructions provided with the PAS staining kit (Beyotime, China). The image data were collected using a DM4B biological microscope (Leica, Bannockburn, IL, USA). The area fraction of collagen or glycogen was calculated as follows: Area fraction of collagen or glycogen (%) = (average collagen or glycogen area/total field area) × 100.

### IHC analysis of tissues

Tissue sections were deparaffinized with xylene and rehydrated through graded ethanol solutions. Antigen retrieval was performed in citrate buffer (pH 6.0), followed by blocking endogenous peroxidase at room temperature. Sections were blocked with 10% normal goat serum for 1 hour, then incubated with primary antibody overnight at 4°C, and secondary HRP - conjugated antibodies for 1 hour at room temperature. Staining was achieved using 3,3’-Diaminobenzidine Tetrahydrochloride (DAB), followed by hematoxylin counterstaining for 3 minutes. Sections were dehydrated with alcohol and cleared with xylene before sealing with neutral gum. Slides were imaged using a DM4B biological microscope (Leica, Bannockburn, IL, USA), and the integrated optical density (IOD) was measured using Fiji/ImageJ software.

### Oil Red O Staining

Oil Red O staining was performed using the ORO kit (PHYGENE, China) according to the manufacturer’s instructions. Staining was inspected and photographed using an inverted microscope (Nikon, Japan). For quantification, the staining was eluted from the cells with isopropanol, and the optical density (OD) was measured at 490 nm using a microplate reader (Molecular Devices), as described before ^63^.

### Statistical analyses

Data are presented as mean±SEM from at least three independent experiments per group, with the exact number indicated in the figures and legends. Statistical analyses of experimental results were conducted using GraphPad Prism v8. A two-tailed Student’s t-test was utilized for comparisons between two groups, while one-way ANOVA followed by Tukey’s multiple post hoc test or Dunnett’s multiple comparisons test was employed to assess differences among multiple groups. A *P* value less than 0.05 was considered statistically significant.

## Acknowledgments

We thank Dr. Ben Chen for performing the orbital adipose tissue sampling.

## Funding

the National Key R&D Program of China 2021YFA1101200 (W.W)

the National Key R&D Program of China 2021YFA0909400 (M.Y.),

the National Natural Science Foundation of China 82171048 (W.W)

the National Natural Science Foundation of China 92253201 (M.Y.)

the National Natural Science Foundation of China 32350026 (M.Y.)

the National Natural Science Foundation of China 22334005 (M.Y.)

the Ministry and the province of Zhejiang Medical and Health Science and Technology Project WKJ-ZJ-2336 (Y.T.)

the Key science and technology program of Wenzhou ZY2022021 (W.Y. and W.W.)

## Author contributions

Examples:

Conceptualization: W.W. and M.Y.

Methodology: Y.Z., E.W., W.L., Z.L., J.W., M.Y., W.W., Y.Z., and E.W.

Investigation: Y.Z., E.W., N.L., L.Z., H.W., S.Y., T.P., Y.H., X.L., X.L., W.Y. and Y.T.

Visualization: Y.Z., W.L.

Funding acquisition: W.W., M.Y. and W.Y.

Project administration: J.W., M.Y. and W.W

Supervision: J.W., M.Y. and W.W

Writing – original draft: Y.Z., J.W. and M.Y.

Writing – review & editing: E.W., W.L., J.W., M.Y. and W.W

## Data and materials availability

All data are available in the main text or the supplementary information.

