## Supplementary information for "TSHR-targeting nucleic acid aptamer treats Graves’ ophthalmopathy via novel allosteric inhibition"

**Table S1. Sequences of synthesized oligonucleotides used in the experiments**

| Sample | Sequence (5’→3’) |
|---|---|
| Library | ACCGACCGTGCTGGACTCANNNNNNNNNNNNNNNNNN NNNNNNNNNNNNNNNNNNNNNNNNNACTATGAGCGAG CCTGGCG |
| YC1 | ACCGACCGTGCTGGACTCAGGGCGCATCTTTGGTAAGCAAA GATGGGTTGCAAGCCTCTCGACTATGAGCGAGCCTGGCG |
| YC2 | ACCGACCGTGCTGGACTCAATCCATCGTCGACGAGCATACGA AGGGAGGGTATTGTAAGGGACTATGAGCGAGCCTGGCG |
| YC3 | ACCGACCGTGCTGGACTCAGATGAACAATCGATGGAAGATT GGGTCGCACCGGCCTCTTTCACTATGAGCGAGCCTGGCG |
| YC4 | ACCGACCGTGCTGGACTCAGGGTGGGTCGTTTCATAGGAAA CGGGTGGAACGCCTCTAGCCACTATGAGCGAGCCTGGCG |
| YC5 | ACCGACCGTGCTGGACTCAGACGGAATCGTCCTTTGTCTACG TGCGGTCTAGACTGTATAGACTATGAGCGAGCCTGGCG |
| YC6 | ACCGACCGTGCTGGACTCAGCCGTGACGTTCTACAGAGAAC GGGTGGATCCCGCCTCTTCACTATGAGCGAGCCTGGCG |
| YC7 | ACCGACCGTGCTGGACTCATGGTCACATCTATCGTACTAGCA TGGGTCAATTGGCCTCTTGACTATGAGCGAGCCTGGCG |
| YC8 | ACCGACCGTGCTGGACTCAGCCGTGACGTTCTACAGAGAAC GGGTGGACCCGCCTCCTCACTATGAGCGAGCCTGGCG |
| YC3-a | ACCGACCGTGCGATGAACAATCGATGGAAGATTGGGTCGCA CCGGCCTCTTTCACTATGAGCGAGCCTGGCG |

| | |
|---|---|
| YC3-b | ACCGACCGTGCTGGACTCAGATGAACTGGGTCGCACCGGCC TCTTTCACTATGAGCGAGCCTGGCG |

**Table S2. Binding free energy decomposition between TSHR and aptamer YC3**

| Residues | Binding free energy (kcal/mol) |
|---|---|
| K146 | -2.67 |
| N170 | -3.18 |
| Q173 | -1.39 |
| T190 | -2.46 |
| Q193 | -2.47 |
| Y195 | -2.67 |
| N198 | -1.87 |
| K218 | -3.51 |
| D219 | -3.46 |
| S243 | -1.06 |
| K244 | -3.23 |
| L265 | -1.23 |
| L266 | -1.36 |
| S268 | -2.94 |

**Table S3. Demographic Information of the Donors Recruited in the Study**

| Age, year/Sex | Duration of GO, year | Smoking Status | Previous Steroid Use | Radiation | Thyroid Treatment |
|---|---|---|---|---|---|
| 33/M | 1 | No | Oral Prednisone | None | No |
| 57/F | 10 | No | Oral Prednisone | None | Methimazole |
| 59/M | 1 | Previous | None | None | Thyroidectomy |
| 50/M | 0.25 | Previous | None | None | Methimazole |
| 36/M | 0.75 | Previous | None | None | Thyroidectomy |
| 55/F | 3 | No | Oral Prednisone | None | Thyroidectomy |
| 20/F | N/A | No | N/A | N/A | N/A |
| 22/F | N/A | No | N/A | N/A | N/A |
| 25/F | N/A | No | N/A | N/A | N/A |
| 24/F | N/A | No | N/A | N/A | N/A |
| 21/F | N/A | No | N/A | N/A | N/A |
| 24/M | N/A | Previous | N/A | N/A | N/A |

F, female; M, male; GO, Graves' ophthalmology; CAS, clinical activity score; N/A, not applicable.

**Table S4. Culture condition and sources of cell lines**

| Cell | Culture condition | Cell Sources |
|---|---|---|
| Human embryonic kidney 293T cells (HEK293T) | high-glucose DMEM (Gibco) containing 10% FBS and 1% penicillin-streptomycin | Procell Life Science & Technology Co. Ltd. (Wuhan, China) |
| Human follicular epithelial cells (Nthy-ori 3-1) | RPMI-1640 medium (Gibco) containing 10% FBS and 1% penicillin-streptomycin | Procell Life Science & Technology Co. Ltd. (Wuhan, China) |
| Human thyroid cancer cells (FTC-133) | RPMI-1640 medium (Gibco) containing 10% FBS and 1% penicillin-streptomycin | Wenzhou Medical University |
| Human thyroid cancer cells (BCPAP) | RPMI-1640 medium (Gibco) containing 10% FBS and 1% penicillin-streptomycin | Wenzhou Medical University |
| Human corneal epithelial cells (HCECs) | high-glucose DMEM (Gibco) containing 10%FBS and 1% penicillin-streptomycin | Wenzhou Medical University |
| Human lens epithelial cells (HLECs) | high-glucose DMEM (Gibco) containing 10% FBS and 1% penicillin-streptomycin | Wenzhou Medical University |
| Human retinal endothelial cells (HRECs) | high-glucose DMEM (Gibco) containing 10% FBS and 1% penicillin-streptomycin | Wenzhou Medical University |
| Adult retinal pigment epithelial cell line-19; (ARPE19) | Dulbecco's Modified Eagle Medium (DMEM), consisting of nutrient Mixture F-12 media | Wenzhou Medical University |

| | | |
|---|---|---|
| | (Gibco) containing 10% FBS and 1% penicillin-streptomycin | |
| The human microglial clone 3 cells (HMC3) | minimum essential medium (Gibco) containing 10% FBS and 1% penicillin-streptomycin | Wenzhou Medical University |

**Table S5. siRNA and random sequences**

| Name | Sequence (5'→3') |
|---|---|
| TSHR siRNA-1 sequences | GTACAACAATGGCTTTACT |
| TSHR siRNA-2 sequences | TGACGTCAATCCCTGTGAA |
| Negative control | UUCUCCGAACGUGUCACGUTT |

**Table S6. Primers used for qPCR**

| Name | Sequence (5'→3') |
|---|---|
| TSHR | F: CCATCAGGAGGAGGACTTCA<br>R: ATTGGGCAGATTAGAAAATG |
| GAPDH | F: AAATCAAGTGGGGCGATGCTG<br>R: GCAGGAGGCATTGCTGATGAT |
| IL-6 | F: GTACATCCTCGACGGCATC<br>R: ACCTCAAACTCCAAAAGACCAG |
| IL-8 | F: GAGAGT GATTGAGAGTGGACC<br>R: ACTGATTCTTGGATACCACAGAG |
| IL-1β | F: GGCTTATTACAGTGGCAATG<br>R: GTAGTGGTGGTCGGAGAT |
| HAS1 | F: TCAAGGCGCTCGGAGATTC<br>R: CTACCCAGTATCGCAGGCT |
| HAS2 | F: GATGACCTACGAAGCGATT<br>R: GCCTGCCACACTTATTGA |

| | |
|---|---|
| HAS3 | F: CGCAGCAACTTCCATGAGG |
| | R: AGTCGCACACCTGGATGTAGT |

F, forward; R, reverse.

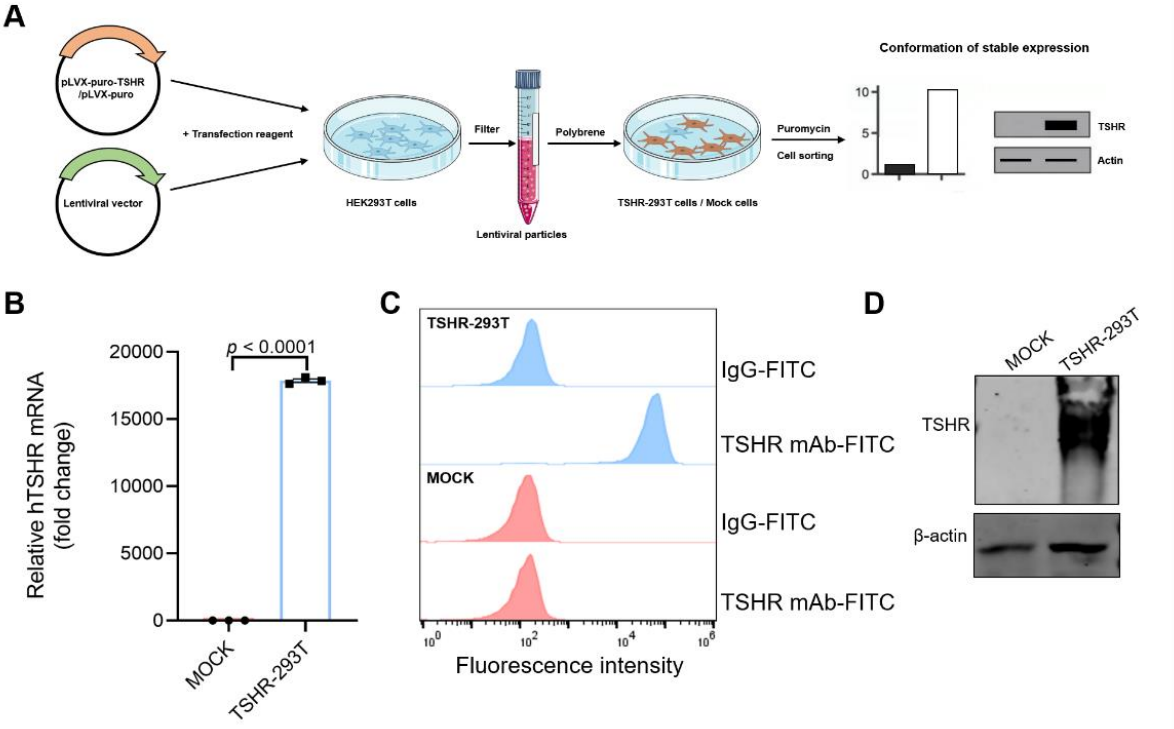


**Fig. S1. Construction and verification of Cell-SELEX model cells.** (**A**) Schematic illustration of the construction of MOCK cells and TSHR-293T cells. (**B**) The relative expression of TSHR mRNA in MOCK cells and TSHR-293T cells was analyzed by qPCR. n=3 independent samples in each group. Data are represented as mean±SEM. Two-tailed Student's t-test was used to calculate *P* values. (**C**) Flow cytometry analysis of the expression of TSHR in MOCK cells and TSHR-293T cells. The FITC-labeled IgG was used as the control. (**D**) Representative immunoblots analysis of TSHR protein level in MOCK cells and TSHR-293T cells. β-actin was used as a loading control.

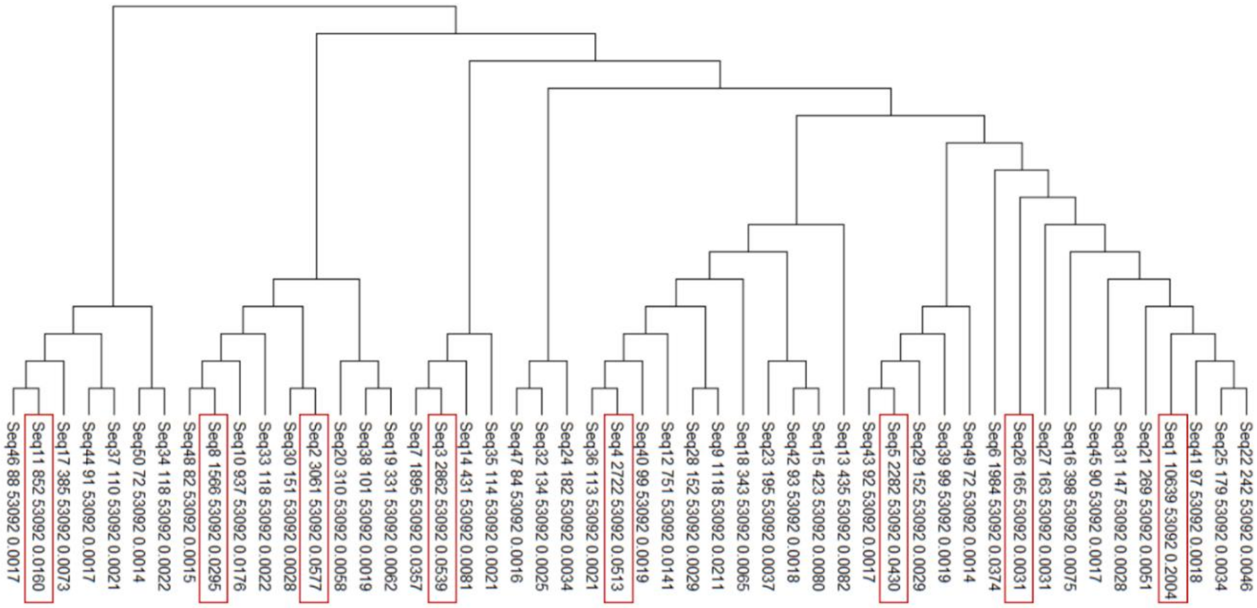


**Fig. S2. Sequence similarity analysis.** Dendrogram of similarity visualization classification (sequence name, number of occurrences, total number of occurrences, and frequency of occurrences from top to bottom) for the 50 individual sequences cloned after the 9th round of screening. Eight representative sequences were selected from different families (boxed).

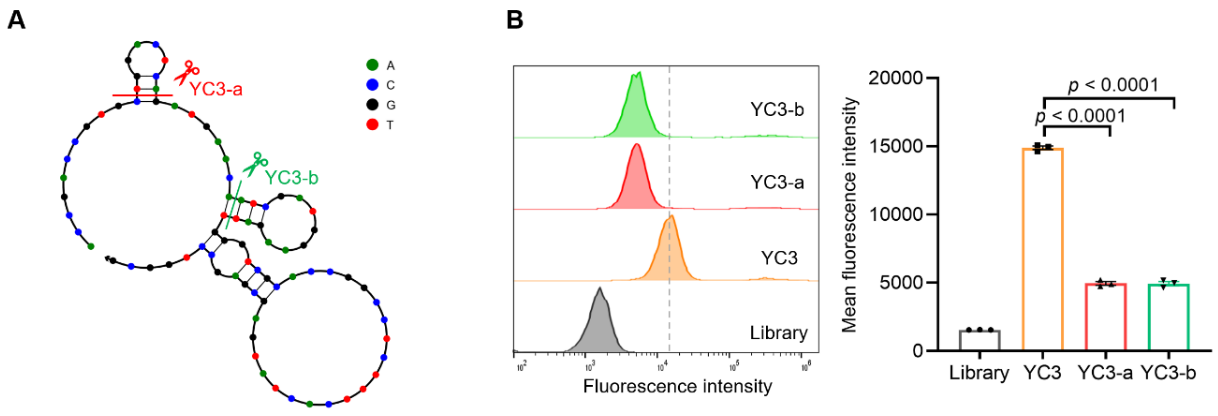


**Fig. S3. Truncation of aptamer YC3.** (**A**) Schematic illustration of two truncated versions of YC3. Secondary structures were predicted by NUPACK software. (**B**) The binding of truncated aptamers (500 nM) to TSHR -293T cells was analyzed by flow cytometry. Quantitative analysis of the relative fluorescence intensity of the aptamers in TSHR-293T cells, n=3 independent samples in each group. Data are represented as mean±SEM. One-way ANOVA, followed by Tukey's multiple post hoc test was used to calculate *P* values.

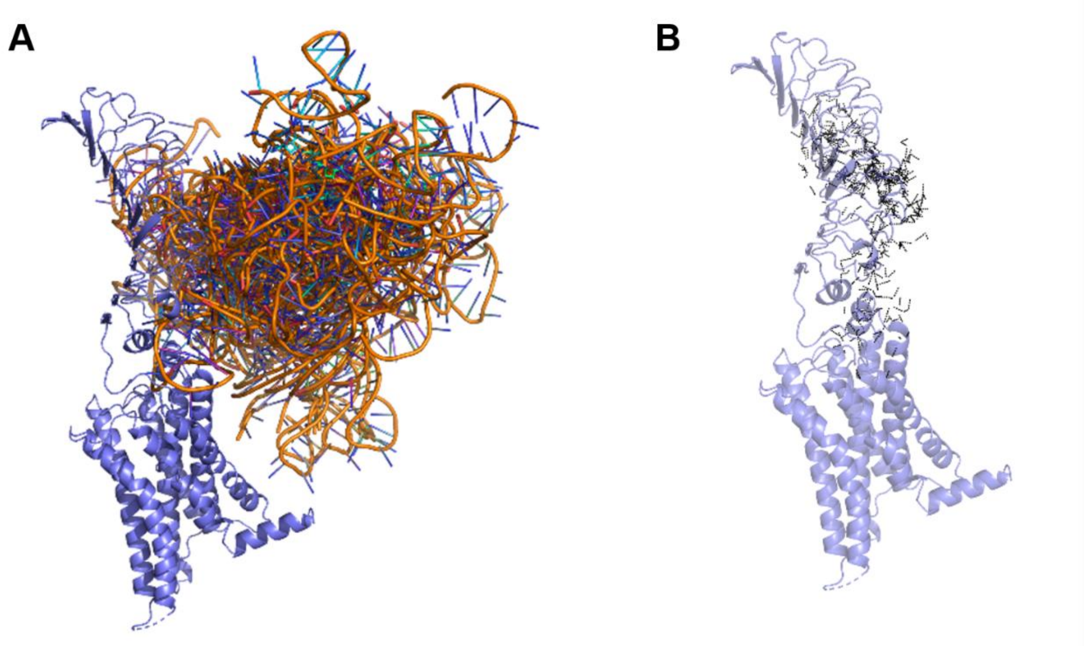


**Fig. S4. Molecular dynamics simulations of TSHR and YC3 docking phases.** (**A**) Molecular dynamic simulation results of *binding sites* between TSHR and the fifty YC3 docking conformations. (**B**) Hydrogen bond network formed between TSHR and all YC3 docking conformations.

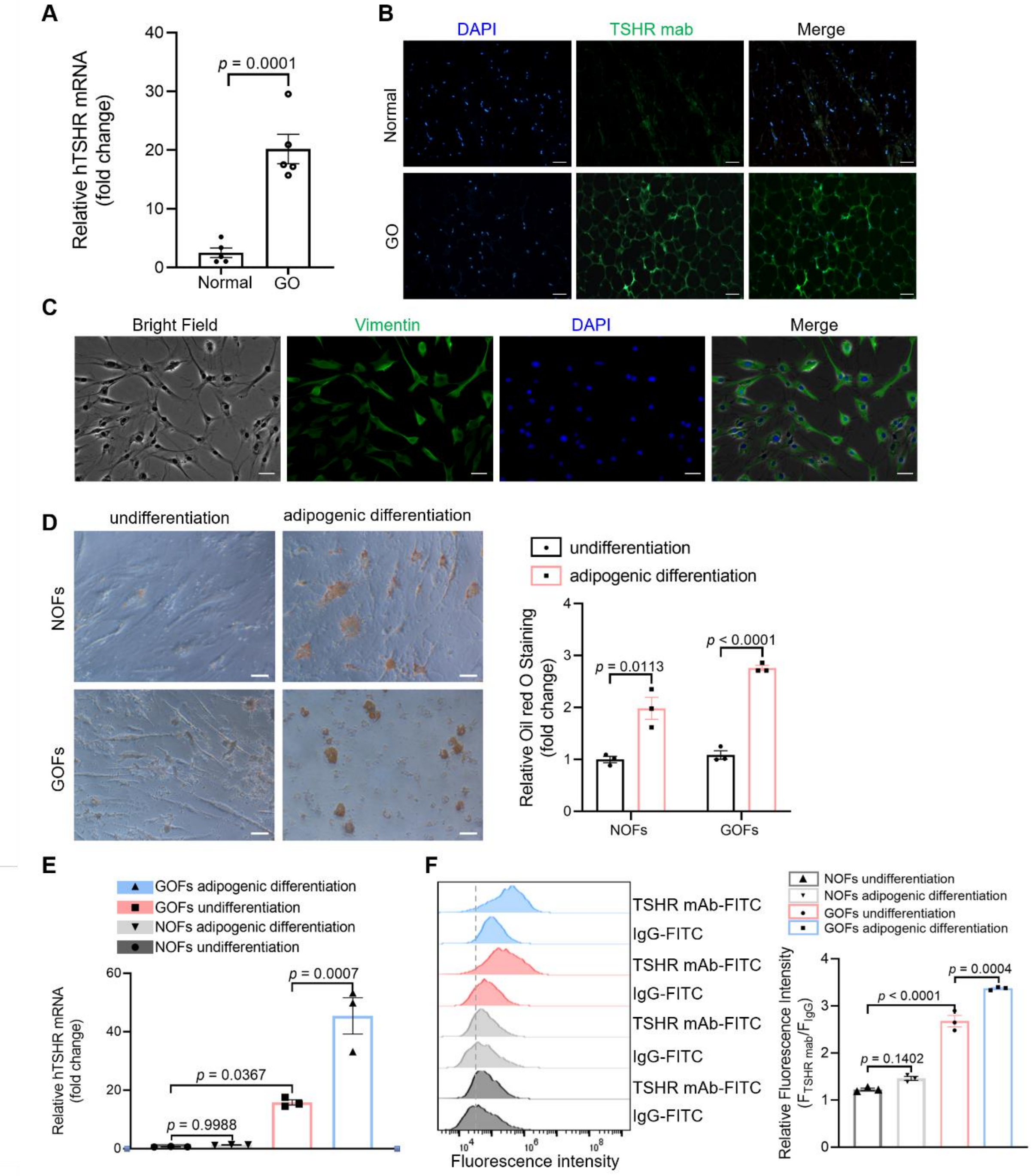


**Fig. S5. TSHR relevance in GO.** (**A**) The mRNA expression levels of TSHR in human orbital adipose tissue from healthy donors or patients with GO, n=5 independent samples in each group. Data are represented as mean±SEM. Two-tailed Student's t-test was used to calculate *P* values. (**B**) Representative images of immunofluorescence staining on human orbital adipose tissue sections from healthy donors or patients with GO for TSHR (green) and DAPI (blue). Scale bars: 50 µm. (**C**) Representative images of immunofluorescence staining on orbital fibroblasts (OFs) from human orbital tissue for Vimentin (green), and DAPI (blue). Scale bars: 50 µm. (**D**) Representative images of Oil Red O staining on GOFs and NOFs, both with and without adipogenic differentiation. Scale bars: 50 µm. Quantitative analysis of the relative Oil Red O staining values of cells, n=3 independent samples in each group. Data are represented as mean±SEM. Two-tailed Student’s t-test was used to calculate *P* values. (**E**) The mRNA expression levels of TSHR in GOFs and NOFs, both with and without adipogenic differentiation, n=3 independent samples in each group. Data are represented as mean±SEM. One-way ANOVA, followed by Tukey's multiple post hoc test was used to calculate *P* values. (**F**) Flow cytometry analysis of the expression of TSHR in GOFs and NOFs, both with and without adipogenic differentiation. Quantitative analysis of the relative fluorescence intensity, n=3 independent samples in each group. Data are represented as mean±SEM. One-way ANOVA, followed by Tukey's multiple post hoc test was used to calculate *P* values.

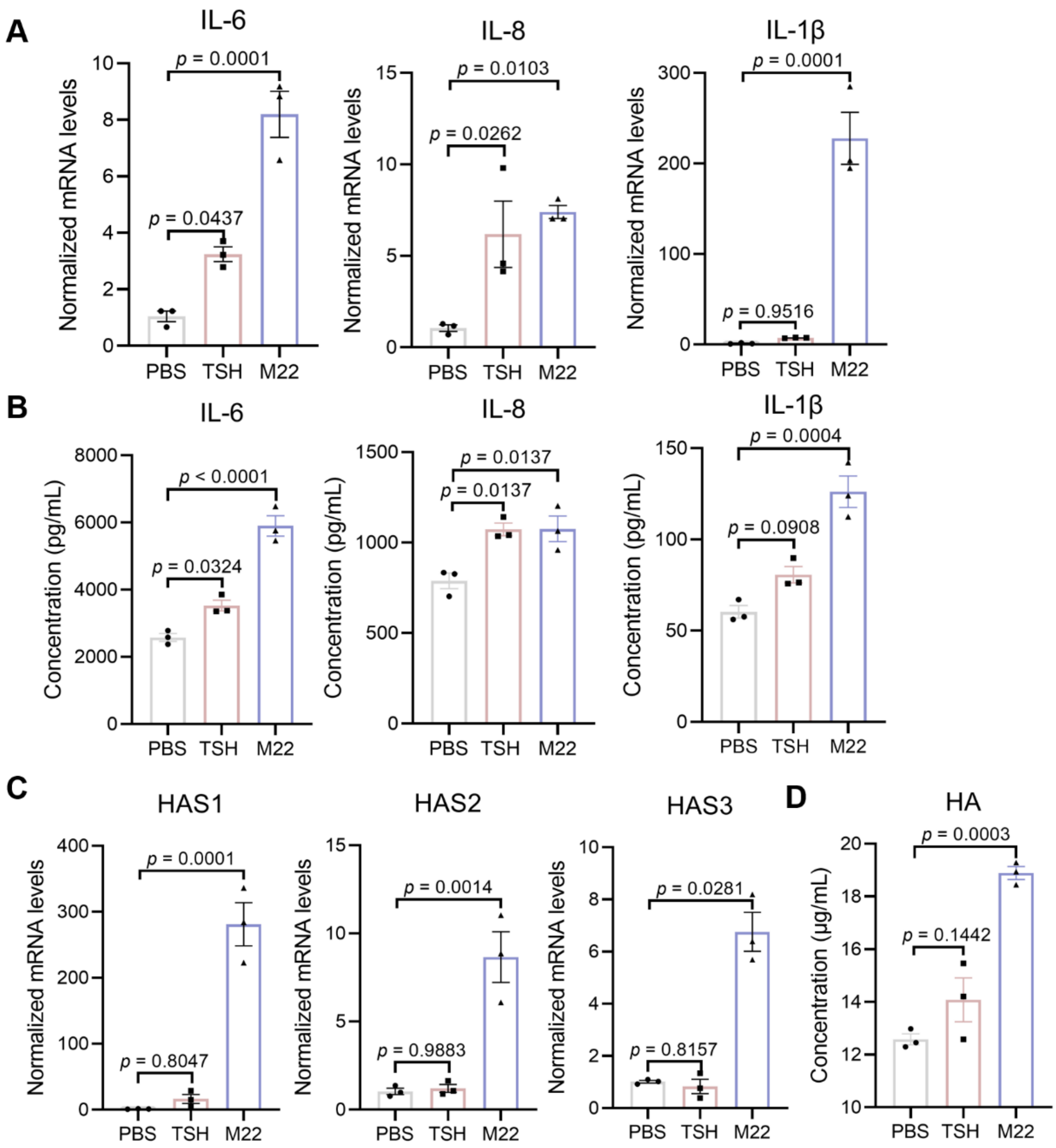


**Fig. S6. Stimulation of TSHR leads to the production of inflammatory cytokines and hyaluronic acid in GOFs.** (**A**) The mRNA expression levels of IL-6, IL-8, and IL-1β in GOFs stimulated by TSH (100 ng/mL) or M22 (100 ng/mL) by qPCR, n=3 independent samples in each group. Data are represented as mean±SEM. One-way ANOVA, followed by Tukey's multiple post hoc test was used to calculate *P* values. (**B**) The concentrations of IL-6, IL-8, and IL-1β in the supernatant medium of GOFs stimulated by TSH (100 ng/mL) or M22 (100 ng/mL) were measured by ELISA, n=3 independent samples in each group. Data are represented as mean±SEM. One-way ANOVA, followed by Tukey's multiple post hoc test was used to calculate *P* values. (**C**) The mRNA expression levels of HAS1, HAS2, and HAS3 in GOFs stimulated by TSH (100 ng/mL) or M22 (100 ng/mL) by qPCR, n=3 independent samples in each group. Data are represented as mean±SEM. One-way ANOVA, followed by Tukey's multiple post hoc test was used to calculate *P* values. (**D**) The concentrations of hyaluronic acid (HA) in the supernatant medium of GOFs stimulated by TSH (100 ng/mL) or M22 (100 ng/mL) were measured by ELISA, n=3 independent samples in each group. Data are represented as mean±SEM. One-way ANOVA, followed by Tukey's multiple post hoc test was used to calculate *P* values.

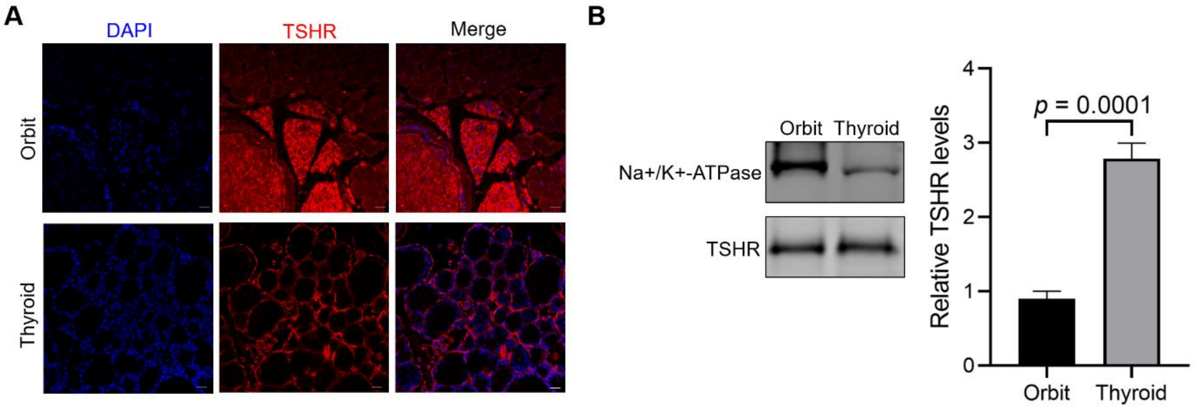


**Fig. S7. TSHR expression in mice thyroidal tissue and orbital tissue.** (**A**) Representative image of immunofluorescence staining on mice thyroidal tissue sections and orbital tissue sections for TSHR (red) and DAPI (blue). Scale bars: 20 μm. (**B**) Representative immunoblots analysis of TSHR protein level in mice thyroidal tissue and orbital tissue. Na+/K+ ATPase was used as a loading control.

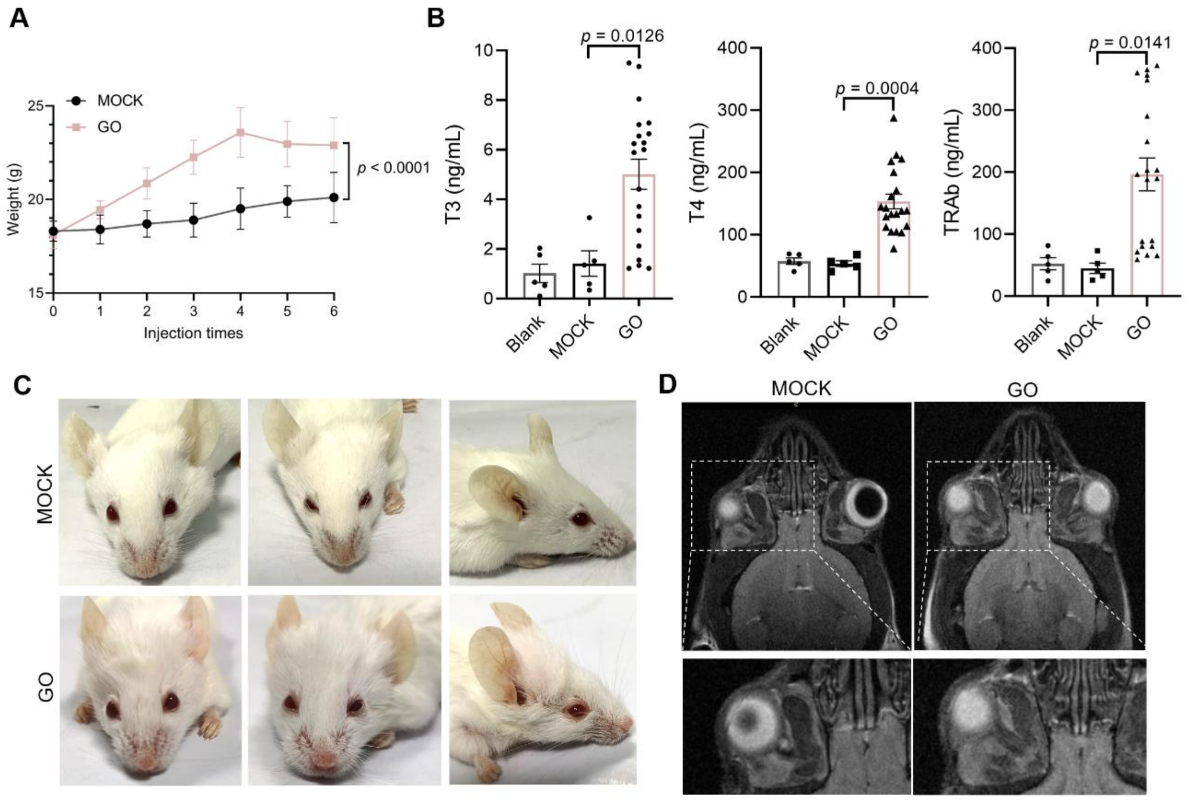


**Fig. S8. Phenotypes of GO mouse model.** (**A**) Measurement of the weight of mice following the injection of Ad-MOCK or Ad-TSHR289 for model construction. Data are represented as mean±SEM. Two-way ANOVA was used to calculate *P* values. (**B**) The concentration of T3, T4, and TRAb in the serum of mice was measured in the Blank group (n=5), MOCK group (n=5), and GO group (n=20) using ELISA. Data are represented as mean±SEM. Two-tailed Student's t-test was used to calculate *P* values. (**C**) Representative images of ocular symptoms in mice from the MOCK group and the GO group. (**D**) Representative images of T2-weighted Magnetic Resonance Imaging of the mice from the MOCK group and the GO group. The lower images provide an enlarged view of the areas within the white dashed boxes shown in the corresponding upper images. White arrows denote the extraocular muscle.

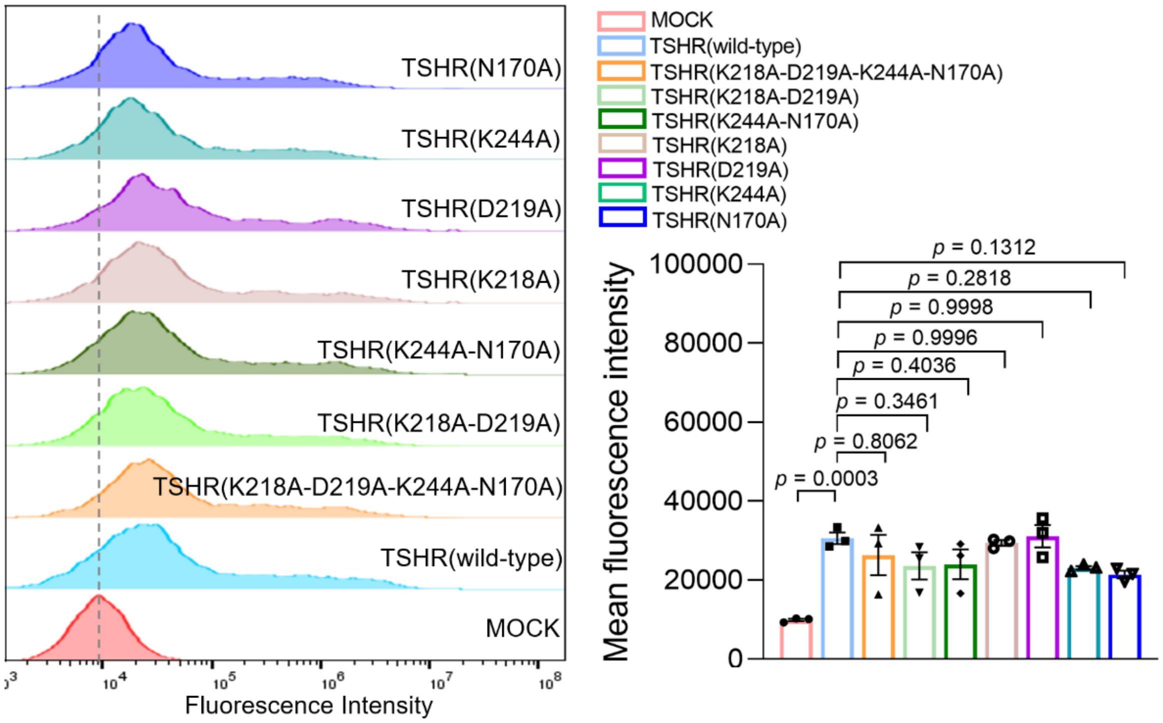


**Fig. S9. Flow cytometry results of TSHR protein expression in HEK293T cells transfected with wild-type and mutant plasmid of TSHR.** Quantitative analysis of the relative fluorescence intensity, n=3 independent samples in each group. Data are represented as mean±SEM. One-way ANOVA, followed by Dunnett's multiple comparisons test was used to calculate *P* values.

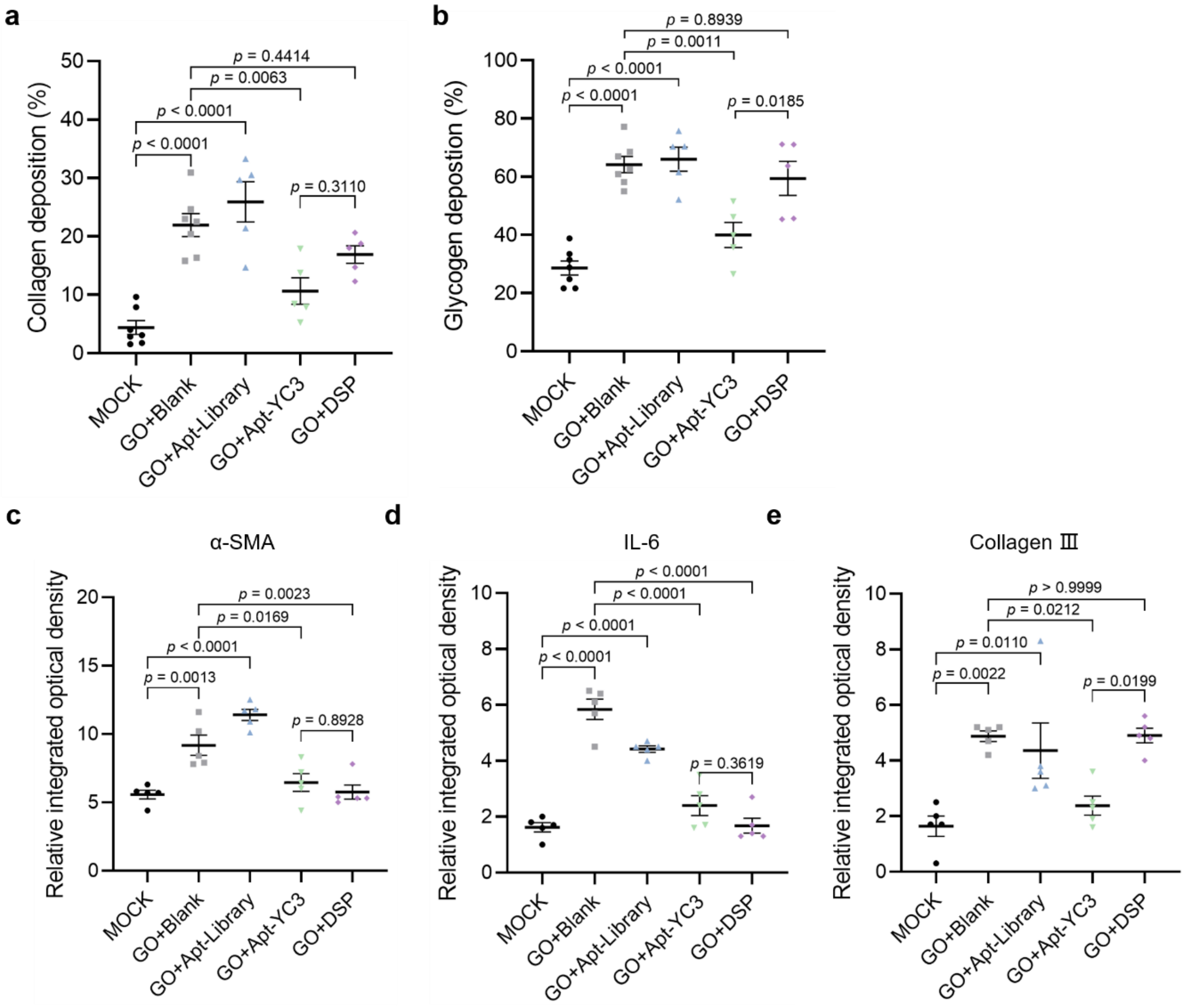


**Fig. S10. Statistical analysis of orbital tissue section staining in each group.** (**A**) Statistical analysis of Masson staining from orbital tissue sections in the MOCK group (n=7), the GO+Blank group (n=7), the GO+Apt-Library group (n=5), the GO+ Apt-YC3 group (n=5), and the GO+DSP group (n=5). Data are represented as mean±SEM. One-way ANOVA, followed by Tukey's multiple post hoc test was used to calculate *P* values. (**B**) Statistical analysis of PAS staining from orbital tissue sections in the MOCK group (n=7), the GO+Blank group (n=7), the GO+Apt-Library group (n=5), the GO+Apt-YC3 group (n=5), and the GO+DSP group (n=5). Data are represented as mean±SEM. One-way ANOVA, followed by Tukey's multiple post hoc test was used to calculate *P* values. (**C** to **E**) Statistical analysis of IHC images by calculating the relative integrated optical density (IOD). Data are represented as mean±SEM (n=5). One-way ANOVA, followed by Tukey's multiple post hoc test was used to calculate *P* values.**{** ADDIN **}**
